# Bacteriocin-mediated prevention of secondary pneumococcal pneumonia by a human commensal *Streptococcus mitis* strain

**DOI:** 10.1101/2025.06.06.658310

**Authors:** Catarina Candeias, João Lança, Marta Alenquer, Daniela Brás, Carina Valente, Maria João Amorim, Raquel Sá-Leão

## Abstract

*Streptococcus pneumoniae* remains a major public health threat despite widespread use of vaccines and antibiotics. Its well-established synergy with influenza A virus (IAV) often results in secondary bacterial pneumonia, a condition associated with high morbidity and mortality. As pneumococcal pneumonia is invariably preceded by nasopharyngeal colonization, preventing colonization represents a promising intervention strategy. Current approaches, including pneumococcal conjugate vaccines and antibiotics, can drive serotype replacement and antimicrobial resistance, underscoring the need for novel, targeted, and serotype-independent alternatives.

We previously identified seven commensal streptococcal strains (*S. oralis* A22 and *S. mitis* B22-G22) that robustly inhibit *S. pneumoniae* growth and biofilm formation in vitro. Here, using a murine model, we demonstrate that colonization with the commensal strain F22Ad significantly reduces pneumococcal density in the nasopharynx by day 10 post-infection and prevents pneumococcal dissemination to the lungs following IAV co-infection. This protective effect was dependent on bacteriocins encoded at the *blp1* locus. Among these, Bac2v1 (alone or in combination with Bac1) significantly reduced pneumococcal colonization. Notably, Bac2v1 exerted serotype-independent inhibitory activity, impairing the growth of diverse pneumococcal strains.

Our findings provide the first in vivo proof-of-concept that commensal streptococci, and specifically their bacteriocins, can disrupt *S. pneumoniae* colonization and prevent progression to secondary bacterial pneumonia. This work introduces a previously unrecognized precision-based approach to pneumococcal disease prevention, offering a potential complement to current strategies.

## Introduction

*Streptococcus pneumoniae* (pneumococcus), a key member of the *Streptococcus* genus, particularly of the streptococci of the mitis group (SMG), is an integral part of the upper respiratory tract (URT) microbiota ^1,2^. While it often colonizes the nasopharynx of humans asymptomatically, it is also a leading cause of significant morbidity and mortality worldwide ^3^. Pneumococcal pneumonia accounts for 25% to 30% of invasive disease in young children, and is the most common clinical presentation of pneumococcal disease in adults ^4^. Additionally, *S. pneumoniae* is a major cause of secondary pneumonia following an influenza A virus (IAV) infection, rendering a mild disease severe or even fatal, particularly in risk groups such as the elderly and immunocompromised individuals ^5-7^. The most widely accepted mechanisms underlying this association involve IAV-induced damage of the respiratory epithelium and neuraminidase-dependent sialic acid release, impaired mucociliary clearance, and weakened innate immune responses, collectively creating an environment permissive to bacterial invasion and proliferation ^8-11^.

Current strategies to prevent and treat pneumococcal infections include multivalent pneumococcal conjugate vaccines (PCVs) and antibiotics, respectively. PCVs target the polysaccharide capsule, the main virulence factor of *S. pneumoniae*, and are effective in the prevention of pneumococcal disease ^12^. Their effectiveness, however, is limited by their valency, as PCVs cover only up to 21 of over 100 pneumococcal capsular types (or serotypes) described ^13,14^. This limited coverage leads to serotype replacement both in colonization and disease ^15,16^. Use of PCVs, together with the pressure exerted by antibiotic use, leads to emergence and dissemination of multidrug-resistant clones expressing serotypes not targeted by current vaccines ^15,17,18^. The limitations of current prevention and treatment options against *S. pneumoniae* demand for development of new therapies, ideally serotype-independent and not prone to emergence of antimicrobial resistance.

The pivotal role of pneumococcal colonization in disease onset, coupled with its high prevalence, underscores its importance and should not be underestimated. *S. pneumoniae* colonizes the nasopharynx in the form of a biofilm – a structured microbial community enclosed in an extracellular matrix and adherent to surfaces or other cells ^19,20^. Colonization is crucial for the evolution of the species, host-to-host transmission and, importantly, it is a prerequisite for disease ^3,21^. Consequently, the World Health Organization has recommended that novel approaches to reduce the burden of pneumococcal disease should target colonization ^21^.

The protective role of commensal bacteria in the URT against invading pathogens – known as colonization resistance - has gained increasing attention ^22,23^. Streptococcal commensals such as *S. salivarius* and *S. oralis* have been studied for their therapeutic and/or prophylactic potential against diseases such as otitis media and pharyngotonsillitis ^24-26^. *S. mitis*, a commensal species phylogenetically related to *S. pneumoniae*, that is generally non-pathogenic, has also been explored for its biotherapeutic potential ^27^. An isolate from the buccal cavity of a human newborn showed significant inhibitory activity against multiple strains of *Haemophilus influenzae*, *Staphylococcus aureus* and *Pseudomonas aeruginosa in vitro*, though its effectiveness under *in vivo* conditions warrants further investigation ^27^. Additionally, *S. mitis* has been investigated for its potential to act as a vaccine, eliciting natural protection against *S. pneumoniae*-induced lung infection through humoral and cell-mediated immune responses ^28,29^. Notably, the strongest protective effect observed so far occurred when *S. mitis* strains, expressing either natural or laboratory-induced *S. pneumoniae* capsules, were used against *S. pneumoniae* of the same serotypes ^28,29^.

Colonization resistance by commensals can occur directly through competition for resources and space or production of antimicrobial compounds such as bacteriocins - ribosomally synthesized antimicrobial peptides that inhibit the growth of closely related or competing bacterial species ^30,31^. The use of bacteriocins as food preservatives and as antimicrobials to inhibit bacterial pathogens has been well-documented ^32^. Notable examples include nisin, produced by *Lactococcus lactis* strains, and lugdunin produced by *Staphylococcus lugdunensis* ^33,34^.

Previously our group identified seven commensal streptococcal strains with *in vitro* inhibitory activity against a diverse collection of *S. pneumoniae* strains covering multiple serotypes and genotypes ^35^. These seven strains (*S. oralis* A22 and *S. mitis* B22 to G22) were found to have a variety of loci encoding putative bacteriocin genes, several of which were absent or extremely rare in *S. pneumoniae*. Deletion mutants indicated that, over half of the bacteriocin loci identified partially or completely explained the anti-pneumococcal activity of the commensal strains. In particular, we observed that for strain F22 deletion of the bacteriocin locus *blp1*, encoding for bacteriocins Bac1 and Bac2, abolished its anti-pneumococcal inhibitory profile. Adding to these results, we have shown that strains A22 to G22 prevented and disrupted pneumococcal biofilms.

The aim of the current study was to assess whether the *in vitro* inhibitory effect observed for these strains was sustained *in vivo*. In particular, we aimed to assess whether strains A22 to G22 could interfere with *S. pneumoniae* colonization as a potential strategy to prevent infection, specifically secondary bacterial pneumonia. To this end, we used a mouse model that combined colonization protocols for *S. pneumoniae* with co-infection by influenza A virus, a well-established trigger of secondary bacterial pneumonia.

Single colonization studies showed that *S. mitis* strain F22 was the best colonizer and, thus, the best candidate for evaluation of its *in vivo* inhibitory activity. We found that by day 10-post IAV infection, strain F22 prevented *S. pneumoniae* D39 from invading the lungs of infected mice by significantly decreasing nasopharyngeal density. Moreover, in the presence of strain F22Δ*blp1*, *S. pneumoniae* D39 invaded the lungs at levels comparable to those found in the control group, highlighting the role of bacteriocins in the observed protective effect. Finally, the potential of bacteriocins encoded by the *blp1* bacteriocin locus to disrupt *in vivo* colonization of *S. pneumoniae* D39 was confirmed using a 3-day treatment regimen, as well as their broad inhibitory effect towards a diverse collection of *S. pneumoniae* serotypes.

We demonstrate the potential of *S. mitis* strain F22 to be used as a biotherapeutic approach to prevent secondary pneumococcal pneumonia, through a mechanism that is dependent on bacteriocin production. These results highlight the versatility and strong potential of bacterial commensals and their bacteriocins as a strategy to reduce pathogen burden.

## Results

### Non-pneumococcal streptococcal strains are able to colonize the nasopharynx of mice

Previously, we identified seven isolates of *S. oralis* (strain A22) and *S. mitis* (strains B22 to G22) species exhibiting inhibitory activity against a diverse collection of *S. pneumoniae* strains^35^. To assess the nasopharyngeal colonization potential of strains A22 to G22, mice were intranasally inoculated with 10^7^ CFU of one of each strain (**Fig. 1a - upper left**). Colonization was evaluated through CFU counts obtained from nasal lavages collected three days post-inoculation. Among the seven strains tested, three (B22, D22 and F22) were detected, albeit at low densities, ranging from 6.7 x 10^1^ to 5.7 x 10^2^ CFU/mL (**Fig. 1b - squares**). To improve colonization, we increased the inoculum concentration to 10^8^ CFU (**Fig. 1a - upper right**). After three days, all strains yielded viable colonies in at least one nasal lavage (range 1-4 mice depending on the strain), with strains B22 and F22 producing higher yields (**Fig. 1b - circles**).

**Fig. 1.**
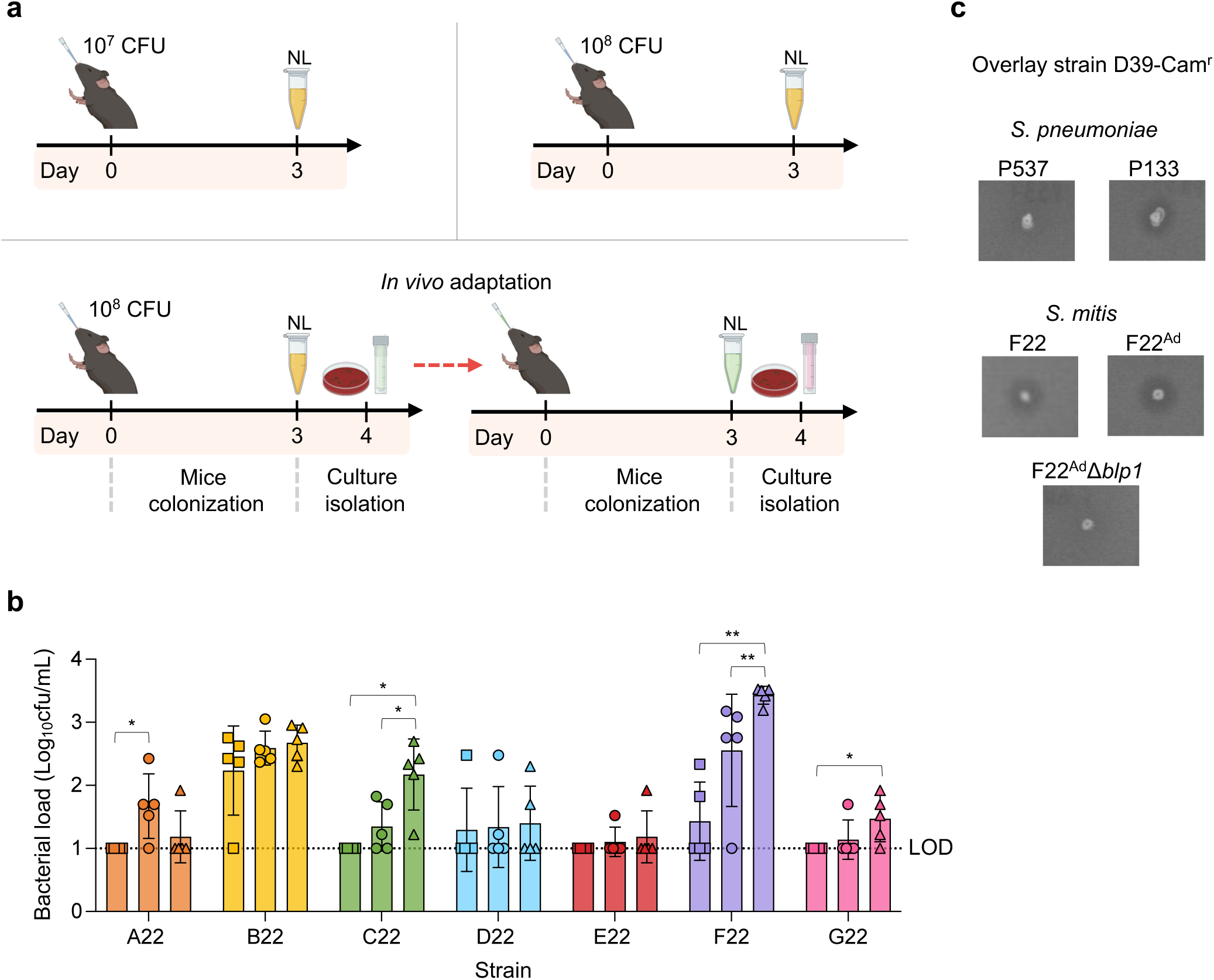
Nasopharyngeal colonization of strains A22 to G22. **a,** Scheme of the experimental design for the nasopharyngeal colonization of strains A22 to G22. Groups of five mice were inoculated with each strain at 10^7^ (upper left) or 10^8^ CFU (upper right) in 10 µL of PBS 1x. After three days mice were sacrificed, and nasal lavages (NL) collected and plated onto GBA. Total bacterial growth was collected and stored in STGG at - 80°C. To adapt strains to mice, samples were sequentially passaged with minimum laboratory manipulations (bottom). NL samples with the highest bacterial loads for each strain were chosen. A loopful of the frozen stock was streaked onto GBA plates and incubated at 37°C in 5% CO_2_. The total bacterial growth of each strain was collected to 200 µL of PBS 1x and 10 µL intranasally inoculated into a new group of five mice. For inoculum evaluation, serial dilutions were plated onto BAP. Mice were sacrificed after three days and this process repeated until the bacterial density stabilized for each strain. Created with BioRender. **b,** Bacterial quantification of NL samples collected from mice intranasally inoculated with strains A22 to G22 at 10^7^ CFU (squares) or 10^8^ CFU (circles) in 10 µL, and sequentially passaged a maximum of four times (last passage in triangles). *p < 0.05, **p < 0.01 (Mann-Whitney U test). The dotted line represents the limit of detection (LOD). **c,** Overlay assays for F22 variants against *S. pneumoniae* D39-Cam^r^. Bacterial growth of each test strain (*S. pneumoniae* strains P537 and P133 as negative and positive control, respectively, and *S. mitis* strains F22, F22^Ad^, F22^Ad^Δ*blp1*), previously inoculated onto BAP, was collected with a 200 µL tip used to stab a TSAgar plate, until the middle of the agar. Stabbed TSAgar plates were incubated for 6h at 37°C in 5% CO_2_. Overlay strain *S. pneumoniae* D39-Cam^r^ was grown in THY until OD_600nm_ of 0.5 was reached and 200 µL added to a tube containing: 5 mL of pre-warmed THY, 4750 U of catalase and 3 mL of molten TSAgar. After 6h of incubation of the TSAgar plate, all content of the overlay strain mixture was gently dispensed on top of the test layer. Plates were left to solidify for 15 min and incubated overnight (without turning over) at 37°C in 5% CO_2_. The next day, plates were examined for the presence of growth inhibition halos, indicative of a positive inhibition result. Each assay was performed in triplicates although only one representative result is shown.

To further improve colonization, we next sought to adapt the commensals to mice by passaging bacteria recovered from the nasal lavages in mice four times using an inoculum of 10^8^ CFU in 10 µL (**Fig. 1a - down**). At this point, the bacterial load/number of colonized mice stabilized for most strains (**Fig. 1b - triangles, and Extended Data Fig. 1**). Bacterial loads obtained for strains C22, F22, and G22 increased significantly by the final passage compared to those obtained in the initial inoculation with 10^7^ CFU. Additionally, this increase was observed not only in the bacterial density but also in the number of mice colonized (range four to five). Strain F22 exhibited the highest bacterial loads (ranging between 1.6 x 10^3^ CFU/mL and 3.4 x 10^3^ CFU/mL) of all strains after sequential passages. Strains D22 and E22 were defined as poor colonizers of the mouse nasopharynx, with a maximum of two out of five mice colonized, and sequential passages failed to increase this number (**Fig. 1b**).

Importantly, inoculation of strains A22 to G22 did not show adverse effects on mouse health, as indicated by stable body weight and the absence of overt clinical signs (**Extended Data Fig. 2**).

Collectively, these results demonstrate that the tested *S. oralis* and *S. mitis* strains (except for strains D22 and E22) can colonize the nasopharynx of mice for three days, particularly with an inoculum of 10^8^ CFU in 10 µL, albeit at low densities. Sequential passages successfully increased the bacterial loads detected, especially for strain F22.

### Sequential passage of *S. mitis* strain F22 in mice results in genetic variants associated with stress response, metabolism, and protein synthesis

Strain F22 was the most effective strain at colonizing the nasopharynx of mice after sequential passages (**Fig. 1b** and **Extended Data Fig. 1**). To identify mutations potentially associated with the increase in bacterial loads following mouse passage, we conducted whole genome sequencing of passaged strains (F22-1, F22-2, F22-3, and F22^Ad^, **Extended Data Fig. 1**). The genome of the original strain F22 was used as reference. After filtering for true variants, we identified a single deletion in strain F22-1, also present in strain F22^Ad^. Additionally, we identified six SNPs exclusively present in strain F22^Ad^ (**Table 1**).

**Table 1.**
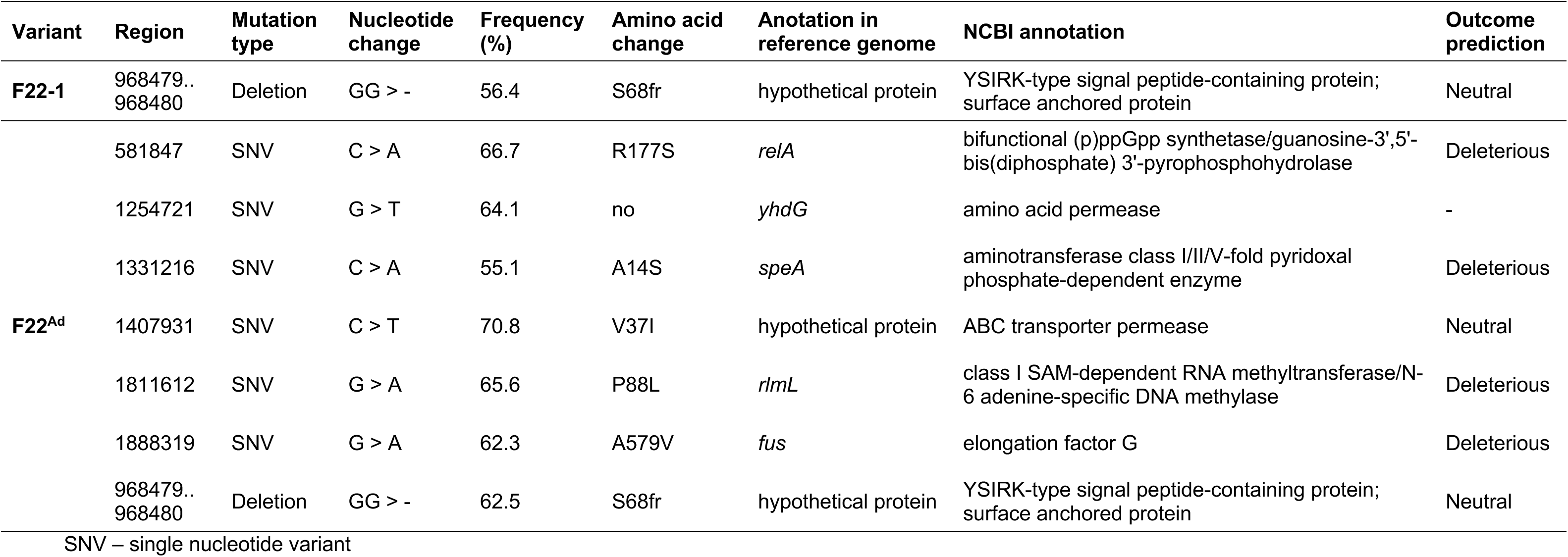
Polymorphisms detected in variants of mouse adapted strain F22.

The deletion found in the genome of strains F22-1 and F22^Ad^ resulted in a frameshift mutation in the terminal region of a YSIRK-type signal peptide-containing protein (identified in a BLASTp search in NCBI). YSIRK-type signal peptides are short amino acid sequences that direct proteins for secretion or transport across the bacterial membrane, many of which are frequently associated with virulence or colonization ^36-39^. A search for this protein in the genome of *S. pneumoniae* TIGR4 revealed homology to the pneumococcal StrH (query cover and identity of 51% and 57%, respectively), a β-N-acetylglucosaminidase virulence factor involved in the degradation of complex N-linked glycans ^40^. Nonetheless, the mutation identified in the present study was predicted to be neutral by PROVEAN, suggesting it is unlikely to significantly affect the function of the protein.

Regarding the additional SNPs observed in strain F22^Ad^, one resulted in a synonymous mutation in *yhdG* gene, which encodes an amino acid permease involved in amino acid transport across cell membranes ^41^. As this mutation did not alter the amino acid sequence of the protein, it should have a neutral effect on its function. In contrast, the remaining mutations introduced non-synonymous changes in *relA*, *speA*, *rlmL* and *fus*, all of which were predicted to be deleterious by PROVEAN, indicating they may impair the proteins’ function. A BLASTp search of these genes in the NCBI database linked them to diverse roles in cellular physiology. Particularly, *relA* is a key player in the stringent stress response, a regulatory mechanism that allows bacteria to adapt to nutrient deprivation and other environmental stresses by altering gene expression, which might have been affected by the identified mutation ^42^. The *speA* gene encodes an enzyme involved in polyamine biosynthesis, a pathway critical for cell growth, stress resistance, and the stabilization of nucleic acids, all of which might have been affected by the identified mutation ^43^. The *rlmL* gene encodes an rRNA methyltransferase involved in ribosomal RNA modification, an essential process for protein synthesis ^44^; the mutation in *rlmL* may imply an effect on ribosome assembly or function, potentially affecting overall protein production. Similarly, the *fus* gene encodes elongation factor G, a key player in the elongation phase of protein synthesis, which is essential for ribosomal translocation along mRNA during translation ^45^. A deleterious mutation in *fus* may disrupt protein elongation, leading to reduced translational efficiency and cellular stress.

Overall, the deleterious mutations identified in *relA*, *speA*, *rlmL*, and *fus* highlight significant functional impacts that could affect the physiology of strain F22^Ad^. These mutations suggest potential adaptations that may influence fitness and competitiveness, particularly in specific environments such as the nasopharynx, where selective pressures could favor such genetic changes.

Importantly, the mouse-adapted strain F22^Ad^ retained the *in vitro* anti-pneumococcal activity displayed by F22 (**Fig. 1c**).

### *S. mitis* prevents the progression of *S. pneumoniae* to the lungs in a mouse model of secondary pneumonia infection following influenza A virus infection, through the action of the *blp1* bacteriocin locus

To evaluate the potential of *S. mitis* to interfere with *S. pneumoniae in vivo*, we selected *S. mitis* strain F22^Ad^ (due to its *in vitro* anti-pneumococcal activity and increased colonization capacity) and *S. pneumoniae* D39-Cam^r^ for proof-of-principle experiments.

First, competition in the nasopharynx was assessed. *S. pneumoniae* and *S. mitis* were inoculated simultaneously or sequentially (**Extended Data Fig. 3a**). Under the conditions tested, *S. mitis* F22^Ad^ did not affect the nasopharyngeal colonization density of *S. pneumoniae* D39-Cam^r^ (**Extended Data Fig. 3b**). Importantly, with this colonization model, we observed that single colonization with *S. pneumoniae* resulted in asymptomatic colonization with no lung infection (**Extended Data Fig.4**).

We next aimed to evaluate whether *S. mitis* could impact *S. pneumoniae* colonization in the context of a previous IAV infection. To this end, mice were intranasally infected with the non-lethal and moderately virulent X31 IAV strain to enable recovery from viral infection. Seven days post IAV infection, different groups of mice were intranasally inoculated with either a mixture of *S. pneumoniae* and *S. mitis* or with each strain alone as controls (**Fig. 2a**). This experimental design is an adaptation of a previously described model for secondary pneumococcal pneumonia ^46,47^. Mice were inoculated 7 days after influenza infection since this timepoint has been associated with the peak of susceptibility to bacterial pneumonia when *S. pneumoniae* is aspirated directly (using between 20 and 50 µL of inoculum) to the lower respiratory tract bypassing nasopharyngeal colonization ^48^. Although this model recapitulates secondary pneumococcal pneumonia, it does not mimic the preceding *S. pneumoniae* colonization period, a fundamental prerequisite for disease in humans ^3^.

**Fig. 2.**
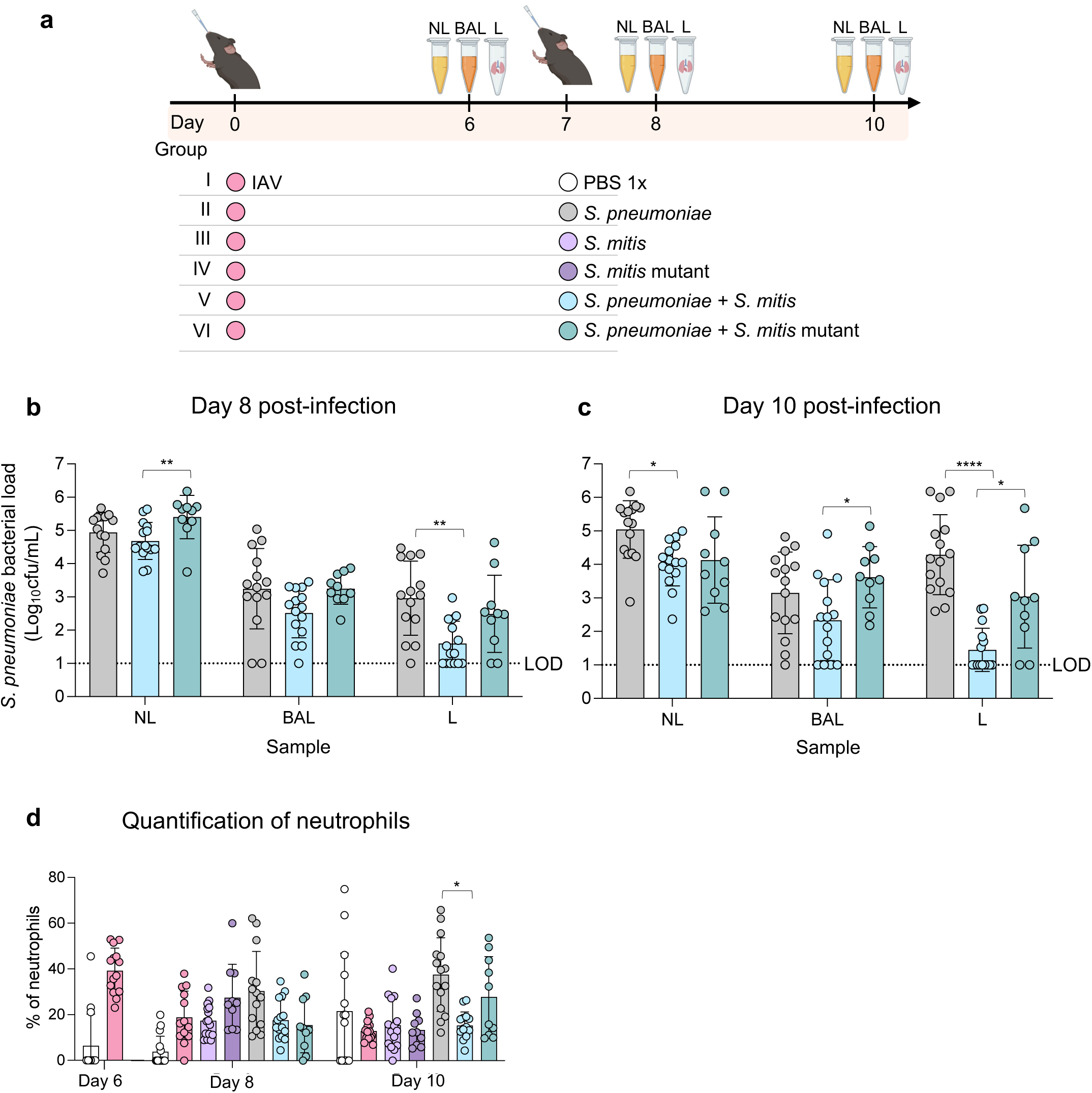
*In vivo* interaction between *S. pneumoniae* and *S. mitis* in the context of an IAV infection. **a,** Experimental design for assessing *S. mitis* impact on colonization and spread of *S. pneumoniae* upon challenge with IAV infection. By day 0, mice were infected with 2000 PFU of IAV X31 in 30 µL of PBS 1x. Seven days after infection, IAV infected mice were separated into groups and inoculated with 10 µL of either *S. pneumoniae* at 10^5^ CFU, *S. mitis* WT at 10^8^ CFU, *S. mitis* mutant at 10^8^ CFU or mixtures of *S. pneumoniae* with each *S. mitis* strain. Six-, eight-, and ten-days post IAV infection, mice were sacrificed, and samples collected: nasal lavages (NL) used for bacterial load assessment; bronchoalveolar lavages (BAL) used for bacterial load assessment and quantification of immune cells by flow cytometry; and lungs (L) used for bacterial load assessment, viral load determination and histological scoring. Created with BioRender. **b, c,** Bacterial loads of *S. pneumoniae* were determined by selective plating serial dilutions of each recovered sample (NL, BAL and L) from mice of all groups sacrificed at days 8 **(b)** or 10 **(c)** post-IAV infection. The dotted line represents the limit of detection (LOD). **d,** Flow cytometry of BAL samples was performed for all groups and the presence of neutrophils analysed. *p < 0.05, **p < 0.01, ****p < 0.0001, ns: non-significant (Kruskal-Wallis followed by Dunn’s multiple comparisons test for multiple comparisons; Mann-Whitney U test to compare two groups)

To overcome this limitation, in our model, we inoculated mice with *S. pneumoniae* in a low volume (10 µL), insufficient for direct inoculation of the lungs, enabling to first establish colonization of the URT. Mice were monitored daily for changes in body weight and development of clinical symptoms. No significant differences were observed in the weight of mice among the various groups, with the maximum weight loss occurring on day 6 post-IAV infection, with approximately 15% weight reduction (**Extended Data Figs. 5a and b**). All mice were able to recover their initial weight. Damage and inflammation of the lungs were evaluated at days 6, 8, and 10 post-IAV infection. The histological score of the lungs was high on day 6 (with an average value of 24 for IAV-infected mice), corresponding to the peak of IAV infection at this timepoint (**Extended Data Fig. 5c**). At days 8 and 10 post-IAV infection, no significant differences in the histological scores were detected between groups (**Extended Data Fig. 5c**). These results collectively support that the severity of lung damage was independent of the bacteria inoculated and, indeed, a result of the IAV infection itself.

The nasopharyngeal loads of *S. pneumoniae*, obtained upon inoculation of mice exposed to IAV infection, were higher (c.a. 2 log) than those observed in the absence of IAV, as previously described (**Figs. 2c and Extended Data Fig.4**) ^49,50^. Particularly, neuraminidases present in both *S. pneumoniae* and IAV are known to be responsible for the desialylation of host cells, releasing high amounts of sialic acid that are used by *S. pneumoniae* ^49,51^. Consequently, *S. pneumoniae* growth in the nasopharynx increases, leading to its aspiration into the lower respiratory tract ^49,51^. In fact, detection of high pneumococcal density in the nasopharynx of humans, associated with a viral infection, has been described as a risk factor for secondary pneumococcal pneumonia ^52,53^. Accordingly, we also detected high densities of *S. pneumoniae* in the bronchoalveolar lavages and lungs at days 8 and 10 post-IAV infection (**Figs. 2b and 2c**), in contrast with what was observed in the absence of IAV (**Extended Data Fig. 4**).

*S. mitis*, by contrast, colonized the nasopharynx of mice at comparable levels regardless of presence or absence of a previous IAV infection, and was rarely detected in the bronchoalveolar lavages or the lungs, both on days 8 and 10 post-IAV infection (**Extended Data Figs. 1, 7a and 7b**).

By day 8 post-IAV infection, no significant differences in bacterial loads of *S. pneumoniae* detected in the nasal lavages were observed between mice inoculated with *S. pneumoniae* alone or in a mixture with *S. mitis* (**Fig. 2b**). However, a significant difference was observed in the bacterial loads recovered from the lungs of mice between the experimental groups. While most mice (13 out of 14) inoculated with *S. pneumoniae* alone presented high bacterial loads in the lungs (ranging from 3.3 x 10^1^ CFU/mL to 2.9 x 10^4^ CFU/mL), six mice inoculated with the mixture of *S. pneumoniae* and *S. mitis* had no *S. pneumoniae* in the lungs, and nine had *S. pneumoniae* at significantly lower densities (2.0 x 10^1^ and 9.3 x 10^2^ CFU/mL, p<0.01).

By day 10 post-IAV infection, we observed a significant difference in bacterial loads of *S. pneumoniae* recovered from the nasal lavages of mice inoculated with *S. pneumoniae* alone or in a mixture with *S. mitis*, indicative of an anti-pneumococcal effect of *S. mitis* on colonization (**Fig. 2c**). Moreover, the preventive effect observed in the lungs by day 8 post-IAV infection was even more pronounced by day 10. While all mice inoculated with *S. pneumoniae* alone presented high bacterial loads in the lungs (ranging from 4.0 x 10^2^ CFU/mL to 1.5 x 10^6^ CFU/mL), nine mice inoculated with the mixture of *S. pneumoniae* and *S. mitis* had no *S. pneumoniae* in the lungs, and six had *S. pneumoniae* at significantly lower densities (3.3 x 10^1^ and 4.8 x 10^2^ CFU/mL, p<0.01).

Taken together, these results suggest that *S. mitis* F22 prevents progression of *S. pneumoniae* D39-Cam^r^ to the lungs in mice previously exposed to an IAV infection, by targeting colonizing *S. pneumoniae*.

Previously in our laboratory, the *in vitro* inhibitory effect of *S. mitis* F22 against *S. pneumoniae* was associated with the presence of the *blp1* bacteriocin locus ^35^. To investigate if this bacteriocin locus was also linked to the prevention of *S. pneumoniae* progression to the lungs following IAV infection, a *blp1* deletion mutant was constructed in the background of *S. mitis* F22^Ad^ (F22^Ad^Δ*blp1*). The loss of anti-pneumococcal inhibitory activity of the adapted mutant *S. mitis in vitro* was confirmed in overlay assays (**Fig. 1c**). Mutant *S. mitis* colonized mice similarly to the WT and it was also rarely detected in the bronchoalveolar lavages or the lungs, both on days 8 and 10 post-IAV infection (**Extended Data Fig. 7b**).

Mice were intranasally inoculated with a mixture of mutant *S. mitis* and *S. pneumoniae*, with the same experimental design described above; as controls, a mixture of WT *S. mitis* and *S. pneumoniae*, or each bacterial strain alone were tested in parallel (**Fig. 2a**).

By day 8 post-IAV infection, *S. pneumoniae* alone or in the presence of mutant *S. mitis* was detected in the lungs of mice at similar densities (**Fig. 2b**). This was in sharp contrast with the lower lung density when *S. pneumoniae* was inoculated in a mixture with WT *S. mitis*. By day 10 post-IAV infection this difference was even more pronounced (**Fig. 2c**). Moreover, mice inoculated with *S. pneumoniae* in the presence of mutant *S. mitis*, were equally colonized in the nasopharynx as mice inoculated with *S. pneumoniae* alone.

Together, these results indicate that the ability of *S. mitis* F22^Ad^ to prevent *S. pneumoniae* migration to the lungs is dependent on the *blp1* bacteriocin locus.

### *S. mitis* prevents lung invasion by *S. pneumoniae* without broad immune cell recruitment

To evaluate the immune response following IAV infection and bacterial challenge, we performed flow cytometry analysis of BAL samples collected on days 6, 8, and 10 post-infection. Single-cell suspensions were stained with a panel of fluorescently labeled antibodies targeting cell surface markers: CD11b and Ly6C for monocytes, GR1/Ly6G for neutrophils, and CD4/CD8 for T cell subsets (**Extended Data Table 3**). Dead cells were excluded using a viability dye, and doublets were removed based on FSC-A/FSC-H gating. Quantification was performed as the percentage of live, CD45+ immune cells within the BAL population for each cell type (**Extended Data Fig. 6)**.

No statistically significant differences were observed in the percentage of monocytes (**Extended Data Fig. 5d**), CD4^+^ (**Extended Data Fig. 5e**) and CD8^+^ cells (**Extended Data Fig. 5f**), across groups challenged with one or more bacterial strains, suggesting that the immune response to IAV was not affected by the presence of bacteria.

Significant differences were observed in the proportion of neutrophils found in the BAL on day 10 post-IAV infection (**Fig. 2d**), which was significantly higher in mice inoculated with *S. pneumoniae* alone and with a mixture of *S. pneumoniae* and mutant *S. mitis*. This finding is in agreement with the higher bacterial loads detected in these groups of BAL samples (albeit statistically non-significant, **Fig. 2c**), compared to its load when inoculated in a mixture with WT *S. mitis*. All together, these results suggest that *S. mitis* F22^Ad^ has a direct preventive effect on the progression of *S. pneumoniae* D39-Cam^r^ to the lungs.

### Bacteriocins from the *blp1* bacteriocin locus decrease colonization by *S. pneumoniae*

To test whether bacteriocins produced by the *blp1* locus of *S. mitis* F22^Ad^ impacted colonization by *S. pneumoniae*, these were first synthesized using a cell-free protein synthesis (CFPS) system ^54^. The *blp1* bacteriocin locus contains two bacteriocins, Bac1 and Bac2, that are not post-translationally modified ^35^. The mature bacteriocin sequences were predicted, based on the conserved amino acid sequence before a double-glycine motif; for Bac2 two putative cleavage sites were found and thus two variants - Bac2v1 and Bac2v2 - were obtained (**Extended Data Table 4**).

To test if these bacteriocins were able to perturb colonization by *S. pneumoniae*, mice were intranasally inoculated with *S. pneumoniae* D39-Cam^r^ at day 0 and colonization established for three days, as described above (**Fig. 3a**). At day three post-inoculation, mice were intranasally inoculated with 10 µL of either Bac1, Bac2v1 or Bac2v2. Moreover, these bacteriocins are predicted to act as a two-peptide due to the presence of GxxxG motifs and thus a mixture of Bac1 and Ba2v1 or Bac1 and Ba2v2 was also intranasally inoculated in a final volume of 10 µL (**Extended Data Fig. 8a**). The CFPS reaction using water instead of DNA was used as negative control. This inoculation was repeated every 12 hours for the following three days. At 12 hours post the last inoculation, mice were sacrificed, NL collected and *S. pneumoniae* bacterial loads quantified.

**Fig. 3.**
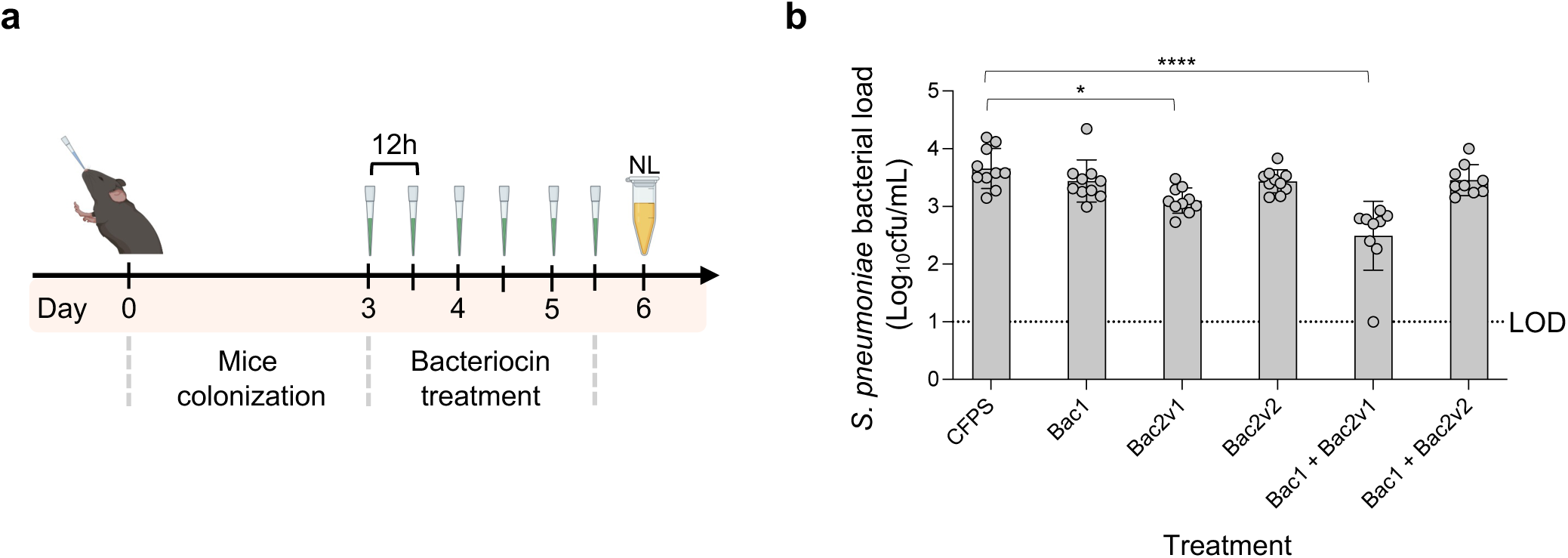
*In vivo e*ffect of *blp1* bacteriocins on *S. pneumoniae* colonization. **a,** Experimental design to evaluate the inhibitory activity of bacteriocins from the *blp1* locus on colonizing *S. pneumoniae*. By day 0, mice were intranasally inoculated with 10 µL of *S. pneumoniae* D39-Cam^r^ at 10^5^ CFU. Three days after bacterial inoculation, 10 µL of Bac1, Bac2v1, Bac2v2 and their combinations were intranasally inoculated every 12h for the following three days. Groups of mice inoculated with the CFPS negative were used as control. Twelve hours after the last bacteriocin inoculation, mice were sacrificed, and nasal lavages (NL) collected. Created with BioRender. **b,** Bacterial loads of *S. pneumoniae* D39-Cam^r^ present in the nasal lavages of each mouse were quantified for the different groups. The dotted line represents the limit of detection (LOD). ***p < 0.001, ****p < 0.0001 (Mann-Whitney U test)

We observed no effect of Bac1 or Bac2v2 alone or in combination when the bacterial loads for *S. pneumoniae* were compared to the CFPS control (**Fig. 3b**). However, a significant decrease in the bacterial loads of *S. pneumoniae* was observed in mice inoculated with Bac2v1. This decrease was even more pronounced when Bac2v1 was inoculated in combination with Bac1 with bacterial loads being approximately 10 times lower than those obtained with the CFPS control (**Fig. 3b**).

Importantly, the three-day treatment with CFPS produced bacteriocins had no adverse effects on mouse health, as evaluated by monitoring mouse weight (**Extended Data Fig. 8b**).

Taken together these results indicate that bacteriocins produced by *blp1* of *S. mitis* F22 decrease *S. pneumoniae colonization in vivo*. The observed activity of Bac2v1 alone suggests it functions independently, while the enhanced effect observed when Bac1 is combined with Bac2v1 may reflect synergistic or additive antimicrobial activity.

### Bac2v1 inhibits epidemiologically relevant *S. pneumoniae*

To assess if the observed inhibitory effect of Bac2v1, alone and in combination with Bac1, was maintained towards a diverse collection of *S. pneumoniae* serotypes and genotypes, a panel of 36 *S. pneumoniae* strains was tested (**Extended Data Table 1**). Planktonic growth was monitored over 24 hours in the presence of either Bac1, Bac2v1, their combination or the CFPS negative control (**Extended Data Fig. 9)**. Bac2v1, either alone or in combination with Bac1, significantly impaired bacterial growth across nearly all tested serotypes. This effect was evident as a reduction in growth rate (**Fig. 4a**) or a delay in the onset of the exponential phase (**Fig. 4b**). Notably, serotypes 14 and 15B/C were the only exceptions, showing no significant response to treatment. Bac1 alone did not exhibit any inhibitory activity.

**Fig. 4.**
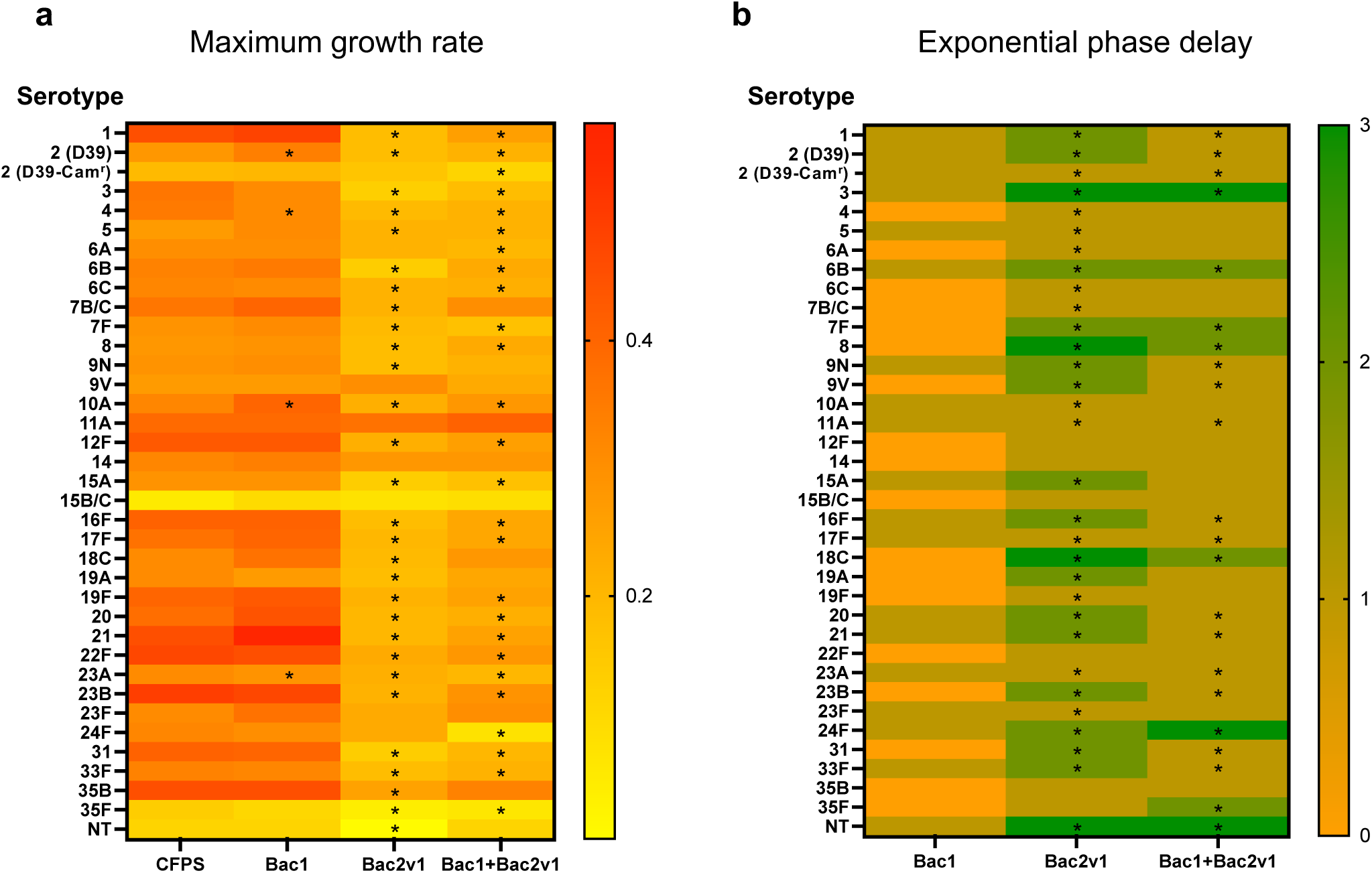
*In vitro e*ffect of *blp1* bacteriocins on *S. pneumoniae* planktonic growth. Cell-free synthesized bacteriocins Bac1 and Bac2v1 were tested against 36 serotypes of *S. pneumoniae.* Pneumococcal cultures were diluted to 10^4^ CFU/mL in fresh C+Y_YB_ medium, distributed in 384-well plates (90 µL per well) and treated with 10 µL of cell-free synthesized bacteriocin (when in single; 5 µL of each when in combination). Controls were treated with the same volume of water, C+Y_YB_ medium or the CFPS reaction using water instead of DNA. Growth was monitored for 24 hours by measuring OD_595nm_ every 30 minutes using a plate reader (Tecan Infinite 200 Pro). Three independent experiments were performed for each bacteriocin. **a,** Maximum growth rate (h^-1^) was calculated for all growth curves, and the mean of three independent replicates is indicated. **b,** The exponential phase delay (in hours), in comparison with the CFPS negative control, was calculated for all treatment groups. The values correspond to intervals of exponential growth delay compared to the control: 0 – timepoint of exponential growth remains equal to or below the control; 1 – timepoint of exponential growth surpasses the control up to 2 hours; 2 – timepoint of exponential growth surpasses the control up to 4 hours; 3 – timepoint of exponential growth surpasses the control more than 4 hours. *p < 0.05 (Mann-Whitney U test)

The inhibitory profile observed *in vivo* for D39-Cam^r^ colonization was recapitulated *in vitro*, with a marked decrease in growth rate upon treatment with Bac2v1 in combination with Bac1 (**Extended Data Fig. 9)**. An enhanced inhibitory effect of the combination, relative to Bac2v1 alone, was also observed for serotypes 6A and 24F. However, for most serotypes, Bac2v1 alone was sufficient to disrupt normal growth dynamics.

These findings demonstrate the broad-spectrum inhibitory potential of Bac2v1 against *S. pneumoniae*, highlighting its promise as a candidate for targeted antimicrobial strategies, particularly when enhanced through combination with Bac1.

## Discussion

Our study demonstrates the potential of a *S. mitis* strain to be used as a live biotherapeutic, particularly as a preventive strategy in the context of a previous IAV infection. Strikingly, inoculation of *S. mitis* strain F22^Ad^ in the nasopharynx of mice during an IAV infection significantly reduced the progression of *S. pneumoniae* to the lungs, a fundamental step for secondary pneumococcal disease. Additionally, this protective effect was dependent on bacteriocins produced by strain F22, which disrupted *S. pneumoniae* colonization in the nasopharynx.

Bacteriocins can also be used for a precision-based alternative or complement to existing interventions. Unlike broad-spectrum antibiotics, which target a wide array of bacteria, or pneumococcal conjugate vaccines, which are limited by their serotype-specificity, these approaches target *S. pneumoniae* directly and independently of its serotype. Both strategies are groundbreaking, representing, to our knowledge, the first demonstration of leveraging a commensal bacterium to prevent secondary pneumonia caused by *S. pneumoniae* following IAV infection. Additionally, the use of a bacteriocin - produced by commensal strains and rarely found within the pneumococcal population - to disrupt *S. pneumoniae* colonization offers a novel and targeted approach to prevent infection.

Previous attempts to demonstrate a protective effect of *S. mitis* against *S. pneumoniae* lung infection in murine models have been limited to a few studies, which employed *S. mitis* strains expressing pneumococcal capsules ^28,29^. While those findings suggested a potential for cross-protection, the serotype-specific nature of the approaches highlighted the need for alternative strategies that confer serotype-independent protection. The protective effect observed in those studies was attributed to both humoral and cell-mediated immune responses, involving antigen-specific IgG and IgA production as well as Th17-mediated mucosal immunity. However, the level of protection was incomplete, as indicated by only a partial reduction in lung bacterial loads, highlighting the limitations of solely modulating the immune system against infections. In the present study, we could not validate a contribution of broad immune cell populations to the protective role of S. *mitis* against S. *pneumoniae* (**Figs. 2d and Extended Data Fig. 5d-f**). The complexity of the mucosal immune landscape and the limited scope of our immune profiling preclude definite conclusions, and this warrants deeper investigations. Nonetheless, the near-complete inhibition of *S. pneumoniae* in the lungs of IAV-infected mice by day 10-post IAV infection, suggests a strong contribution from non-immune mechanisms.

Instead, the absence of *S. pneumoniae* in the lungs of mice was dependent on bacteriocins, particularly of the *blp1* locus (**Figs. 2b and c**). Our findings open the possibility of using bacteriocins in the control of bacterial infections. Other studies have reported the inhibitory role of *blp*-encoded bacteriocins in *S. pneumoniae,* where the regulatory mechanism of the *blp* locus closely resembles the *blp1* locus identified in strain F22 ^35,55^. For example, BlpM and BlpN produced by *S. pneumoniae* were shown to inhibit growth of immunity-deficient pneumococcal strains during colonization, confirming their role in intraspecies competition *in vivo* ^56^. Similarly, BlpI, BlpJ and BlpK demonstrated potent anti-pneumococcal inhibition in biofilms: BlpI and BlpJ as a two-peptide bacteriocin and BlpK alone ^57^.

Unlike such studies that focused on the role of bacteriocins in mediating intraspecies competition within *S. pneumoniae*, our work demonstrates the critical role of *S. mitis* bacteriocins in interspecies inhibition of *S. pneumoniae*. Notably, Bac1 and Bac2 were absent in a collection of 7,548 *S. pneumoniae* genomes ^35^, underscoring their potential for therapeutic application. Furthermore, antimicrobial peptides of diverse origin such as cecropin P1, indolicidin, peptide glycine-leucine amide, pleurocidin, and tachyplesin II were previously demonstrated to be less prone to develop resistance than antibiotics ^54,58^. Future studies using experimental evolution approaches will evaluate whether this is the case for Bac2v1, as well as for the combination of Bac1 and Bac2v1. These studies will be conducted either independently or in conjunction with antibiotics targeting *S. pneumoniae*, to assess potential synergistic or complementary effects.

In conclusion, our study highlights *S. mitis* and its bacteriocins as promising tools to target *S. pneumoniae* colonization and prevent secondary pneumococcal pneumonia upon IAV infection. By exploring the natural ecological interactions between commensal and pathogenic species, we present a targeted, serotype-independent strategy that overcomes the limitations of current interventions. Moving forward, future research should focus on the application of *S. mitis* as a live biotherapeutic in human clinical trials, which would be a key step in translating these findings into practical healthcare applications to transform the prevention and treatment of pneumococcal diseases in the near future.

## Methods

### Strains used in this study

The strains used in this study are described in **Extended Data Table 1**. This included *S. oralis* A22, *S. mitis* B22 to G22, *S. pneumoniae* D39, *S. pneumoniae* D39-Cam^r^, a collection of *S. pneumoniae* expressing 35 different serotypes, and influenza A virus A/HKx31 (X31). *S. oralis* strain A22 and *S. mitis* strains B22 to G22, recently described, were isolated from human nasopharyngeal samples and, *in vitro*, have the capacity to inhibit a diverse collection of *S. pneumoniae* ^35^. *S. pneumoniae* strain D39-Cam^r^ is a chloramphenicol resistant variant of the laboratory strain D39 and was used as a representative of its species for all *in vivo* experiments ^59,60^. Strain influenza A virus X31 contains six segments of the mouse adapted A/Puerto Rico/8/34 (PR8), and segments 4 and 6 (that express hemagglutinin (Ha or H) and neuraminidase (Na or N), respectively) from the H3N2 influenza A/Hong Kong/1/68, rendering it less virulent and adequate for experiments requiring the survival for influenza infected mice for posterior inoculation with streptococci ^61,62^.

### Ethics Statement

This study was ethically reviewed and approved by both the Ethics Committee and the Animal Welfare Body of the Instituto Gulbenkian de Ciência (IGC) (license reference A009/2019), and the Direção Geral de Alimentação e Veterinária (DGAV - license reference 0421/000/000/2020), the Portuguese National Entity that regulates the use of laboratory animals. All experiments followed the Portuguese (Decreto-Lei n° 113/2013) and European (Directive 2010/63/EU) legislations, concerning animal welfare, housing, and husbandry.

### Mice infection and tissue collection

All experiments were performed using six to eight-week-old littermate C57BL/6J female mice under specific pathogen-free conditions at the IGC biosafety level 2 (BSL-2) animal facility. A one-week adaptation period to the BSL-2 facility was completed for all animals. Animals were group housed in individually ventilated cages with access to food and water *ad libitum*. Littermates were randomly allocated to experimental conditions in groups of five unless otherwise stated. Intranasal inoculations were performed while mice were under general anesthesia with inhaled isoflurane (Abbot), as previously described ^46^. Briefly, mice were held in the palm of the hand, upright, and the bacterial/viral suspension was gently administered into the nostrils using a micropipette. Mice were daily monitored for clinical symptoms such as weight loss, lethargy, and ruffled fur. To comply with the best animal welfare practices, animals that presented a reduction of initial bodyweight of more than 25% were sacrificed. At indicated timepoints, animals were sacrificed with CO_2_ and nasal lavage (NL), bronchoalveolar lavage (BAL) and lungs were collected.

NL was collected as previously described ^63^. Briefly, after sterilization of the fur with 70% ethanol, a longitudinal cut along the midline of the neck of the animal was performed to make the trachea accessible. Once the trachea was properly exposed, a small cut about half-way up was performed. The trachea was cannulated towards the nose, and 1 mL of sterilized PBS 1x dispensed. The NL was collected to a 1.5 mL tube placed beneath the nose of the animal and immediately put on ice.

BAL of the whole lung was collected as previously described ^64^. Taking advantage of the small cut performed in the trachea for NL collection, a new catheter was inserted towards the lungs, and 1 mL of sterilized PBS 1x flushed and collected with a 1 mL syringe. The lungs were exposed for confirmation of inflation and absence of leakage.

Lungs were collected for different assays: (i) for histology, the left lobe was collected in one piece to 10% buffered formalin; (ii) for plaque assays, the right lower lobe was collected to a 1.5 mL tube and placed on dry ice; (iii) for bacterial loads detection, the remaining right lobe was collected to an Eppendorf tube and placed on ice.

### Colonization model for bacterial strains

The colonization model used was based on what has been previously described for *S. pneumoniae* (**Fig. 1a**) ^46^. Strains were grown in THY until mid-exponential phase (OD_600nm_ of 0.5) and inoculated intranasally at 10^7^ or 10^8^ colony forming units (CFUs) in 10 µL of sterilized PBS 1x. Serial dilutions were plated onto tryptic soy agar plates supplemented with 5% sheep blood (BAP) (BBL^TM^ Stacker^TM^ Plates, BD) for inoculum concentration confirmation. Plates were incubated overnight at 37°C in 5% CO_2_ and CFUs were counted. Three days after inoculation, mice were sacrificed and NL collected. For bacterial quantification, 10-fold serial dilutions of each murine NL sample were plated in BAP supplemented with 5 µg/mL of gentamicin (GBA) to prevent growth of contaminants. After overnight incubation at 37°C in 5% CO_2_, CFUs were enumerated. In parallel, 100 µL of each nasal lavage were streaked onto GBA plates and, after overnight incubation at 37°C in 5% CO_2_, all visible bacterial growth was collected to 500 µL of skim milk-tryptone-glucose-glycerol (STGG) medium ^65^, and stored at - 80°C. Glycerol was added to NL to a final concentration of 15% before storage at -80°C.

### Mouse adaptation of non-pneumococcal streptococcal strains A22 to G22

*In vivo* adaptation of strains A22 to G22 to the nasopharynx of mice was performed with some modifications to what was previously described for *S. pneumoniae* (**Fig. 1a**) ^66^. Briefly, mice were inoculated with each strain at 10^8^ CFU in 10 µL of PBS 1x, sacrificed after three days, and NL collected and processed as detailed above. Thereafter, total bacterial growth samples stored in STGG from the mice colonized with the highest bacterial load of each strain were chosen. A loopful of the frozen stock was streaked onto GBA plates and incubated at 37°C in 5% CO_2_. On the following day, for each strain, all bacterial growth was collected to 200 µL of PBS 1x, centrifuged at 9600 g for 10 min, and the pellet resuspended in 100 µL of PBS 1x. Ten µL of this bacterial suspension were intranasally inoculated in each mouse. Inoculum concentration was evaluated as described above. Mice were sacrificed after three days and the process was repeated until stabilization of the bacterial density obtained three days post mouse inoculation was reached.

### Strain confirmation by PCR

The presence of the inoculated strains in samples recovered from mice was confirmed by PCR. For detection of *S. pneumoniae* strain D39-Cam^r^, previously described primers targeting the capsular polymerase gene *wzy* of serotype 2 were used (**Extended Data Table 2)**. For detection of strains A22 to G22, we took advantage of previously described genes known to be present in commensal *Streptococcus* species and absent in *S. pneumoniae* (SM12261_1076 and SM12261_1300) and designed specific primers for each strain (**Extended Data Table 2)** ^67^. PCR was performed in a final volume of 10 μL with GoTaq Flexi Buffer 1x, MgCl_2_ 2.5 mM, dNTPs 0.05 mM, primer forward 0.4 μM, primer reverse 0.4 μM and GoTaq 0.025 U/μL. The PCR program used was: initial denaturation at 95°C for 2 min; 30 cycles of denaturation at 95°C for 30 s, annealing at 55°C for 30 s and extension at 72°C for 15 s; final extension at 72°C for 5 min and stop at 16°C.

### DNA extraction of mouse adapted variants of strain F22

Samples with the highest bacterial loads obtained from each successive passage of strain F22 were selected (**Extended Data Fig. 1**). Frozen stocks were plated onto GBA and incubated overnight at 37°C under a 5% CO_2_ atmosphere. On the next day, 500 μL of sterile PBS 1x was added to each plate, cultures resuspended, and the bacterial content collected. From this bacterial suspension, 200 μL were transferred into a tube already containing 200 μL of lysis buffer (MagNA Pure Compact Nucleic Acid Isolation Kit, Roche Diagnostics GmbH) and 17.4 μL of RNase A (100 mg/mL) and incubated for 20 min at 37°C. Total DNA was extracted using the MagNA Pure Compact Nucleic Acid Isolation Kit (Roche Diagnostics GmbH) following the manufacturer’s instructions. The quality of DNA was assessed by measuring A_260_/A_280_ and A_260_/A_230_ ratios using Nanodrop (ThermoFisher Scientific). DNA quantity was measured using the dsDNA High Sensitivity Qubit kit (ThermoFisher Scientific) as per the manufacturer’s protocol. Finally, DNA integrity was evaluated by performing electrophoresis on a 1% agarose gel.

### Whole genome sequencing

Whole genome sequencing (WGS) was performed at the Genomics Unit of IGC, Oeiras, Portugal. Genomes of samples were sequenced on Illumina NextSeq 500 with a target coverage of 100x. Quality control and trimming of paired-end reads was performed using CLC Genomics Workbench version 9.5.1 software (Qiagen, Venlo, The Netherlands), from this point forward called CLC Genomics Workbench for simplicity.

### Variant calling

Reads of each F22 mouse-adapted samples were mapped against the genome of parental F22 strain (accession number ERS19845131). Single nucleotide polymorphisms (SNPs) and small insertions and deletions (InDels) were detected using the Fixed Ploidy Variant Detection tool of the CLC Genomics Workbench software, with the ploidy set to 1 and the variant probability to 90%. To obtain a list of all variants detected, minimum values of coverage (1x), read count (n=2) and frequency (1%) were set. A true variant was defined based on parameters adapted from Lieberman *et al*. ^68^: i) at least 15 reads in each direction, ii) at least 100 bp distance from the reference contig boundaries, iii) average base quality ≥ 20, iv) average mapping quality ≥ 34, and v) p ≥ 0.05 supporting a null hypothesis that the frequency in both read directions is similar. A minimum value for frequency in the population was not set because sequenced samples were not pure (DNA was extracted directly from the bacterial growth of nasal lavages). CLC Genomics Workbench was also used to identify amino acid changes of true SNPs (using the Amino Acid Changes tool). The impact of amino acid changes on gene function was assessed using the Protein Variation Effect Analyzer (PROVEAN) software, which calculates a similarity score based on protein alignment ^69^. Scores below - 2.5 indicate a deleterious effect, while those above this threshold suggest a neutral effect.

### Construction of F22^Ad^Δ*blp1* mutant

F22^Ad^Δ*blp1* mutant was generated by allelic replacement of the bacteriocin-immunity region of *blp1* with a kanamycin resistance gene (*kanR*) flanked by *lox66* and *lox71* sites as previously described ^35^. For that, two fragments flanking the bacteriocin-immunity region of the *blp1* locus were amplified from genomic DNA and the *lox66-P3-KanR-lox71* cassette was amplified from pKan (synthesized by TwistBioscience). Fragments were purified using the Zymoclean™ Gel DNA Recovery Kit (Zymo Research) and ligated by Gibson Assembly (NEB). Nested primers were used to amplify the desired product. Primers used to construct the fragments for transformation are listed in **Extended Data Table 2**. For transformation, strains were grown in C+Y_YB_ medium ^70,71^ without shaking at 37°C until an OD_600_ of 0.5. Cultures were diluted 1:100 in fresh C+Y_YB_ and grown until an OD_600nm_ of 0.04. At that point, cultures were supplemented with 150 ng/ml of DNA and 300 nM of cognate competence stimulating peptide (CSP) (for strain F22 the CSP sequence is NH2-ESRVSRIILDFLFLRKK-COOH ^35^, obtained from NZYTech, >85% purity) and further incubated for 4h. Transformants were selected in BAP supplemented 1 mg/mL of kanamycin (Kan) and confirmed by PCR.

### Overlay assays

Overlay assays were performed as previously described ^72^. Briefly, a loopful of each test strain (F22, F22^Ad^, F22^Ad^Δ*blp1*, *S. pneumoniae* strain P537 as negative control and *S. pneumoniae* strain P133 as positive control ^72^) frozen pre-inoculum was inoculated onto BAP and incubated overnight at 37°C in 5% CO_2_. Tryptic soy broth (Bacto^TM^, BD) with 1.5% agar (Fisher Scientific) (TSAgar) plates supplemented with 4750 U of catalase (Sigma-Aldrich) in a total volume of 20 mL were prepared. On the following day, the bacterial growth of each test strain was collected from the BAP with a 200 µL tip, which was used to stab the TSAgar plate, until approximately half of the agar deepness. Stabbed TSAgar plates were incubated for 6h at 37°C in 5% CO_2_. Overlay strain *S. pneumoniae* D39-Cam^r^ was grown in THY until OD_600nm_ of 0.5 was reached. Two hundred µL of this culture were added to a tube containing 5 mL of pre-warmed THY, 4750 U of catalase and 3 mL of molten TSAgar. After 6h of incubation of the TSAgar plates, all content of the overlay strain mixture was gently dispensed on top of the test layer. Plates were left to solidify for 15 min and incubated overnight (without turning them upside down) at 37°C in 5% CO_2_. On the next day, plates were examined for the presence of growth inhibition halos around the stabbed test strains, indicative of a positive inhibition result.

### *In vivo* interaction between F22^Ad^ and D39-Cam^r^

The *in vivo* inhibitory activity of *S. mitis* strain F22^Ad^ was tested against *S. pneumoniae* D39-Cam^r^ (**Fig. 2a**). Mice were inoculated with a mixture of *S. pneumoniae* at 10^7^ CFU/mL and *S. mitis* at 10^10^ CFU/mL in 10 µL. A 1000x higher inoculum of *S. mitis* was necessary to ensure that similar bacterial loads were obtained after three days of colonization for both species; these inoculum concentrations were used throughout the study. Inoculation with each strain alone was done as control. For inoculum concentration confirmation, serial dilutions were plated onto BAP and BAP supplemented with 4 µg/mL of chloramphenicol for strain differentiation. Mice were sacrificed after three days post inoculation and NL collected. Bacterial quantification was performed by 10-fold serial dilution of each NL sample, plated in parallel in GBA and GBA supplemented with 4 µg/mL of chloramphenicol for strain differentiation. In parallel, 100 µL of each nasal lavage was streaked onto GBA plates and, after overnight incubation at 37°C in 5% CO_2_, all bacterial growth was collected to 500 µL of STGG medium, and stored at -80°C. Glycerol was added to NL to a final concentration of 15% before storage at -80°C.

The inhibitory activity of *S. mitis* against *S. pneumoniae* was also tested at other time points (**Fig. 2a**). Mice were inoculated with either *S. pneumoniae* or *S. mitis*. Three days afterwards mice were inoculated with the other species (*S. mitis* or *S. pneumoniae*, respectively). For comparison, control mice were inoculated with PBS 1x instead of the competing strain. Mice were sacrificed three days after the last inoculation, and NL collected. Bacterial quantification was performed as detailed above.

### *In vivo* interaction of F22^Ad^, and its *blp1*-deficient mutant with D39-Cam^r^ in the context of a IAV infection

The *in vivo* inhibitory activity of adapted *S. mitis* strains WT (F22^Ad^) and mutant (F22^Ad^Δ*blp1*) (inoculated at 10^10^ CFU/mL) was tested against *S. pneumoniae* D39-Cam^r^ in the context of a IAV infection (**Fig. 3a**). Mice were infected with 2000 plaque forming units (PFU) of X31 in 30 µL of PBS 1x. Six days after infection, IAV and mock-infected mice were sacrificed for viral load assessment. Seven days after infection, IAV infected mice were separated into groups inoculated with 10 µL of either *S. pneumoniae*, *S. mitis* WT, *S. mitis* mutant, or each *S. pneumoniae*/*S. mitis* mixtures. Groups of mice inoculated with PBS 1x were used as control. For inoculum concentration confirmation, serial dilutions were plated onto BAP and BAP supplemented with 4 µg/mL of chloramphenicol for strain differentiation. Mice were sacrificed one or three days after bacterial inoculation (8- or 10-days post IAV infection, respectively) and NL, BAL and lungs were collected. Bacterial quantification was performed as detailed above for all sample types.

### Plaque assays

Viral loads were determined at day 6 post-infection from the right lower lobes of IAV and mock-infected mice. Lung weight was calculated and, for each mg of lung, 10 µL of serum free media was added. Samples were homogenized using tungsten carbide beads (Qiagen) in a TissueLyser II (Qiagen) at 20 s^-1^ for 3 min and supernatants collected. Titration by plaque assay was performed as previously described using Madin-Darby Canine Kidney cells ^73,74^.

### Histology

Histological scoring was performed as in Nieto *et al.* ^75^. Briefly, left lung lobes were collected and fixed in 10% buffered formalin for 24 hours, embedded in paraffin, divided into longitudinal sections (3 μm thick) and stained with hematoxylin and eosin. Histological scoring was performed blindly by a pathologist of the Histology Facility of IGC. The score quantitatively measures the extent of tissue damage, with higher values indicating increasingly severe pathological changes.

### Flow cytometry

The presence of immune cells in the BAL of infected mice was analyzed by flow cytometry. Samples were centrifuged and transferred to a V-bottom 96-well plate (Thermo Scientific). Fc blocking (rat anti-mouse CD16/CD32, clone 2.4G2, BD Pharmingen^TM^) was performed to minimize unspecific staining, and cells were incubated 20 min at 4°C. Primary antibodies were added according to **Extended Data Table 3** and cells were incubated 25 min at 4°C, in the dark with agitation. Live/Dead Fixable Yellow Dead Cell Stain Kit (Invitrogen^TM^) was used, and cells were incubated 20 min at room temperature in the dark. Finally, cells were fixed with Intracellular (IC) Fixation Buffer (Invitrogen^TM^) according to manufacturer’s recommendations. Flow cytometry analysis of cell populations was performed in a BD LSR Fortessa X-20 SORP (BD Biosciences) equipped with BD FACSDiva^TM^ 8 (BD Biosciences) and FlowJo^TM^ software v10.10 (BD Life Sciences).

Cell populations were first gated by forward and side scatter to exclude debris, followed by doublet discrimination using FSC-A versus FSC-H. Live single cells were then gated as CD45⁺, and immune subsets c of specific markers: monocytes (CD11b⁺ Ly6C⁺), neutrophils (CD11b⁺ GR1/Ly6G⁺), CD4⁺ T cells (CD3⁺ CD4⁺), and CD8⁺ T cells (CD3⁺ CD8⁺) (**Extended Data Fig. 6)**.

### Cell-free protein synthesis (CFPS) of bacteriocins

To identify which bacteriocins of the *blp1* locus of *S. mitis* strain F22 (Bac1, Bac2v1 or Bac2v2)^35^ have anti-pneumococcal activity, optimized bacteriocin-gene fragments were designed to achieve maximal expression. The mature bacteriocin sequences were predicted based on conserved amino acid sequence before the double-glycine motif ^76^. Corresponding mature nucleotide sequences were codon optimized for *Escherichia coli* (**Extended Data Table 4**). Additionally, for each bacteriocin, the ribosome-biding site (RBS) was optimized to maximize the translation rate in *E. coli* (**Extended Data Table 4**). In the end, DNA fragments were designed with minimal T7 promoter (TAATACGACTCACTATAG), RBS, initiation codon (ATG), mature bacteriocin gene codon optimized for *E. coli*, a spacer sequence (GCTTTATCTGAGAATATACTCGAA), and T7 terminator (CCCCTAGCCCGCTCTTATCGGGCGGCTAGGGG) (**Extended Data Table 4**). Then, CFPS of bacteriocins was performed using PUREfrex®2.0 (GeneFrontier Corporation, Japan) as previously described ^54^. For each bacteriocin, a final concentration of 10 nM of respective DNA fragment was used. As control, a reaction with the same volume of water (instead of DNA) was made. All reactions were incubated for 4 hours at 37°C.

### *In vivo* effect of bacteriocins on *S. pneumoniae* colonization

The *in vivo* inhibitory activity of Bac1, Bac2v1, Bac2v2 and their combinations was tested against *S. pneumoniae* D39-Cam^r^ in the mouse model of pneumococcal nasopharyngeal colonization (**Fig. 5a**) ^46^. Briefly, mice were intranasally inoculated with 10^5^ CFU in 10 µL of *S. pneumoniae*. Three-day colonized mice were then intranasally inoculated with 10 µL of each cell-free synthesized bacteriocin, alone or in combination, every 12 hours for the following three days. The CFPS reaction using water instead of DNA was used as control. Mice were sacrificed 12 hours after the last inoculation, NL collected, and bacterial quantification performed as described above.

### *In vitro* effect of bacteriocins on *S. pneumoniae* planktonic growth

Cell-free synthesized bacteriocins were tested against 36 serotypes of *S. pneumoniae* (**Extended Data Table 1**). Pneumococcal cultures were grown in C+Y_YB_ medium at 37°C until reaching an OD_600nm_ of 0.5. These cultures were then diluted to 10^4^ CFU/mL in fresh C+Y_YB_ medium, distributed in 384-well plates (90 µL per well) and treated with 10 µL of cell-free synthesized bacteriocin (when in single; 5 µL of each when in combination). Controls were treated with the same volume of water, C+Y_YB_ medium or the CFPS reaction using water instead of DNA. Growth was monitored for 24 hours by measuring OD_595nm_ every 30 minutes using a plate reader (Tecan Infinite 200 Pro). Three independent experiments were performed for each bacteriocin.

### Statistical analysis

GraphPad Prism version 9.5.0 for macOS (GraphPad Software, Boston, Massachusetts, USA) was used for all statistical analyses. The Kruskall-Wallis test followed by Dunn’s multiple comparisons test was used for multiple comparisons. The Mann-Whitney U test was used for comparisons between two groups. Statistical significance was represented as *p<0.05, **p<0.01, ***p<0.001, ****p<0.0001.

## Acknowledgments

The authors are grateful to the Animal House Facility, Flow Cytometry Facility, Histopathology Unit and Genomics Unit of Instituto Gulbenkian Ciência for technical support, sample processing and data collection. We also acknowledge Sérgio Filipe (FCT NOVA, Portugal) and Marc Veldhoen (iMM, Portugal) for helpful discussions, Ana Cristina Paulo (ITQB NOVA) for help with the statistical analyses, and Amir Pandi for the support in the implementation of the CFPS for bacteriocin synthesis.

## Funding

The work developed in the Sá-Leão laboratory was funded by FCT - Fundação para a Ciência e a Tecnologia, I.P., through project STOPneumo (PTDC/BIA-MIC/30703/2017), MOSTMICRO-ITQB R&D Unit (UIDB/04612/2020, UIDP/04612/2020), and LS4FUTURE Associated Laboratory (LA/P/0087/2020). CC, JL and DB were supported by grants PD/BD/148434/2019, UI/BD/153385/2022 and PD/BD/148391/2019, respectively. The work developed in the Amorim laboratory was funded by FCT and the “La Caixa” Foundation under the grant agreement HR22-00722. Universidade NOVA de Lisboa has filed a provisional patent application that covers pharmaceutical compositions comprising strains A22 to G22 and/or bacteriocin molecules, and derivatives thereof, which can inhibit the growth and persistence of the pathogen *S. pneumoniae* (PT119647).

## Notes

### Competing Interest Statement

The authors have declared no competing interest.

