## Supplemental Tables for "Bacteriocin-mediated prevention of secondary pneumococcal pneumonia by a human commensal *Streptococcus mitis* strain"

**Extended Data Table 1. Strains used in this study**

| Strain | Species | Observations | Reference |
|---|---|---|---|
| A22 | *S. oralis* | Commensal strain isolated from the URT of an adult ≥ 60 years old | 1 |
| B22 | *S. mitis* | Commensal strain isolated from the URT of a child ≤ 6 years old | 1 |
| C22 | *S. mitis* | Commensal strain isolated from the URT of a child ≤ 6 years old | 2 |
| D22 | *S. mitis* | Commensal strain isolated from the URT of an adult ≥ 60 years old | 3 |
| E22 | *S. mitis* | Commensal strain isolated from the URT of a child ≤ 6 years old | 1 |
| F22 | *S. mitis* | Commensal strain isolated from the URT of a child ≤ 6 years old | 1 |
| $F22^{Ad}$ | *S. mitis* | Mouse adapted variant of F22 | This study |
| $F22^{Ad}$Δ*blp1* | *S. mitis* | Mouse adapted variant of F22 with a deleted *blp1* bacteriocin locus | This study |
| G22 | *S. mitis* | Commensal strain isolated from the URT of a child ≤ 6 years old | 1 |
| X31 | Influenza A virus | Mildly virulent H3N2 virus | 4,5 |
| D39-$Cam^r$ | *S. pneumoniae* | Serotype 2 laboratory reference strain | 6,7 |
| P537 | *S. pneumoniae* | Serotype 6A bacteriocin-naïve strain (negative control) | 8 |
| P133 | *S. pneumoniae* | Serotype 6A bacteriocin-producer strain (positive control) | 8 |
| PT9520 | *S. pneumoniae* | Serotype 1 ST306 | 1 |
| PT12491a | *S. pneumoniae* | Serotype 3 ST180 | 9 |
| PT3392 | *S. pneumoniae* | Serotype 4 | 10 |
| PT6718 | *S. pneumoniae* | Serotype 5 ST1223 | 1 |
| PT11898 | *S. pneumoniae* | Serotype 6A ST65 | 9 |
| PT12762 | *S. pneumoniae* | Serotype 6B ST469 | 9 |
| PT12587 | *S. pneumoniae* | Serotype 6C ST386 | 9 |
| EL2307 | *S. pneumoniae* | Serotype 7B/C ST1201 | 3 |
| PT8227 | *S. pneumoniae* | Serotype 7F ST191 | 1 |
| PT12584 | *S. pneumoniae* | Serotype 8 ST53 | 9 |
| PT12470 | *S. pneumoniae* | Serotype 9N ST66 | 9 |
| PT3386 | *S. pneumoniae* | Serotype 9V | 10 |
| PT12656 | *S. pneumoniae* | Serotype 10A ST1551 | 9 |
| PT12509 | *S. pneumoniae* | Serotype 11A ST62 | 9 |
| PT10316 | *S. pneumoniae* | Serotype 12F ST989 | 1 |

(continues next page)

**Extended Data Table 1. (cont.)**

| Strain | Species | Observations | Reference |
|---|---|---|---|
| PT12593 | *S. pneumoniae* | Serotype 14 ST156 | 9 |
| PT12598 | *S. pneumoniae* | Serotype 15A ST14367 | 9 |
| PT12545 | *S. pneumoniae* | Serotype 15B/C ST1262 | 9 |
| PT12592 | *S. pneumoniae* | Serotype 16F ST30 | 9 |
| PT10418 | *S. pneumoniae* | Serotype 17F ST123 | 1 |
| PT10457 | *S. pneumoniae* | Serotype 18C ST13409 | 1 |
| PT12645 | *S. pneumoniae* | Serotype 19A ST416 | 9 |
| PT12574 | *S. pneumoniae* | Serotype 19F ST9972 | 9 |
| PT10682 | *S. pneumoniae* | Serotype 20 ST1026 | 1 |
| PT12537 | *S. pneumoniae* | Serotype 21 ST432 | 9 |
| PT12329 | *S. pneumoniae* | Serotype 22F ST10220 | 9 |
| PT12515 | *S. pneumoniae* | Serotype 23A ST42 | 9 |
| PT12507 | *S. pneumoniae* | Serotype 23B ST8959 | 9 |
| PT12719 | *S. pneumoniae* | Serotype 23F ST338 | 9 |
| PT12770a | *S. pneumoniae* | Serotype 24F ST16 | 9 |
| PT12492 | *S. pneumoniae* | Serotype 31 ST1766 | 9 |
| PT12052 | *S. pneumoniae* | Serotype 33F ST9607 | 9 |
| PT12558 | *S. pneumoniae* | Serotype 35B ST198 | 9 |
| PT12430 | *S. pneumoniae* | Serotype 35F ST1635 | 9 |
| PT12401 | *S. pneumoniae* | Non-typeable ST3097 | 9 |

**Extended Data Table 2. Primers used in this study**

| Purpose | Name | Sequence | Description |
|---|---|---|---|
| Amplification of serotype 2 *wzy*[a] | F_serotype 2 | TATCCCAGTTCAATATTTCTCCACTACACC | Confirmation of *S. pneumoniae* D39-Cam[r] |
| | R_serotype 2 | ACACAAAATATAGGCAGAGAGAGACTACT | |
| Amplification of locus no. SM12261_1076[b] | F1_1076 | ATTATGGGACCGCCATGTCATC | Confirmation of strains A22, C22 and D22 |
| | F2_1076 | ATTATGGGACCGTCATGTCATC | Confirmation of strains B22, E22, F22 and G22 |
| | R1_1076 | GTCAAGTCGGGCCGTCTCAATC | Confirmation of strain A22 |
| | R2_1076 | GTCAAGTCTGGCCGTCTCAATC | Confirmation of strains B22 and C22 |
| | R3_1076 | GTCAAATCTGGTCGTCTCAATC | Confirmation of strain D22 |
| | R4_1076 | GTCAAGTCTGGTCGTCTCAATC | Confirmation of strains E22, F22 and G22 |
| Amplification of locus no. SM12261_1300[b] | F1_1300 | AATCGCCATGTTCAGGTGATGG | Confirmation of strain A22 |
| | F2_1300 | AATCGTCATGTGCAGGTTATGG | Confirmation of strains B22, C22 and E22 |
| | F3_1300 | AATCGTCACGTGCAGGTTATGG | Confirmation of strains D22, F22 and G22 |
| | R1_1300 | ATGATCGAAGGTCCTGTAAGGC | Confirmation of strain A22 |
| | R2_1300 | ACAATGGAAGGACCTGTCAGGC | Confirmation of strain B22 |
| | R3_1300 | CAATGGAAGGACCTGTTAGGC | Confirmation of strains C22, D22, E22, F22 and G22 |
| Deletion of *blp1* from strain F22 | P1F_blp1_F22_new | TGTTAAAGGTAATCAGCTGACC | Upstream *blp1* locus of strain F22 |
| | P1R_ blp1_F22_new | **GTCAATACCGTTCGTATAGCATACATTATACGAAGTTAT**GCAACCTTGACAACATC | |
| | P2F_ blp1_F22_pB2.6 | **GATATATTGATGTTGTCAAGGTTGC**ATAACTTCGTATAATGTATGC | Lox-Kan-Lox from plasmid pB2.6 |
| | P2R_ blp1_F22_pB2.6 | **GATAACCACTTAAACACTAAGTACAC**ATAACTTCGTATAGCATAC | |
| | P3F_ blp1_F22_new | **GTTTTAGTACCGTTCGTATAATGTATGCTATACGAAGTTAT**GTGTACTTAGTGTTTAAGTGG | Downstream *blp1* locus of strain F22 |
| | P3R_ blp1_F22_new | AGTTTTCAGCAGTAATTCCTGG | |
| | P4F_ blp1_F22 | GTATTGTTTTCTAAGCGCTGCG | Internal primers for final construct |
| | P4R_ blp1_F22 | GGGCTTGAATACTTGCTTGG | |

[a]From https://www.cdc.gov/strep-lab/media/pcr-oligonucleotide-primers.pdf
[b]From reference [11].
**Bold:** Homology sequences for final fragment assembly

### Extended Data Table 3. Antibodies used for flow cytometry

| Antibody | Target | Brand | Catalog | Clone | Host | Diluted 1: |
|---|---|---|---|---|---|---|
| CD3-FITC | Mouse | BioLegend | 100204 | 17A2 | Rat | 100 |
| CD4-BV421 | Mouse | BioLegend | 100438 | GK1.5 | Rat | 100 |
| CD45.2-APC | Mouse | BioLegend | 103112 | 30-F11 | Rat | 100 |
| CD8-BV711 | Mouse | BioLegend | 100748 | 53-6.7 | Rat | 100 |
| GR1/Ly-6G-PE | Mouse | BD Pharmingen | 551461 | 1A8 | Rat | 200 |
| Ly-6C-PerCPCy5.5 | Mouse | eBioscience | 45-5932-80 | HK1.4 | Rat | 200 |
| CD11b/Mac1-BV605 | Mouse | IGC Antibody Facility | - | M1/70 | Rat | 100 |

### Extended Data Table 4. DNA fragments optimized for cell-free protein synthesis

| Bacteriocin | Ribosome-binding site (RBS) | Mature nucleotide sequence codon optimized for *E. coli* (including ATG) |
|---|---|---|
| Bac1 | CATTAATCTGGCGTTAG ACGGCAGGACTACGAC ATTAAGGAGGTTTTTT | ATGGACAAGGTGGGCGCGGGCGAGGTGGTGCAGGCGCTGGG CATCTGCACCATCGGCGGCGCGGCGCTGGGCAGCGTGATCC CGGTGGTGGGCACCCTGGCGGGCGGCGTGCTGGGCGCGCA GTTCTGCACCGCGGCGTGGGGCGCGTTCCGTGCGAGCTAA |
| Bac2v1 | AACAACAAGTAATTGAG TATCCTAATAGAGGAG GTATTTTA | ATGGACAAGCAGCTGGCGGACCGTTTCCTGAGCGGCGTGGG CGGCGCGGTGGAGGGCCTGAGCCTGTGCGCGCAGACCATCC CGTTCCCGACCCCGCAGATCTACCTGATCTGCGCAGCAGGAG GAGCAGCAGCAAGTGTTCTGTGGCCGCATTAA |
| Bac2v2 | CTAGTTCAATAAACTTA TTAAGGAGGTATTTT | ATGGCGGTGGAGGGCCTGAGCCTGTGCGCGCAGACCATCCC GTTCCCGACCCCGCAGATCTACCTGATCTGCGCAGCAGGAGG AGCAGCAGCAAGCGTGCTGTGGCCGCATTAA |
