## Supplemental Figures for "Bacteriocin-mediated prevention of secondary pneumococcal pneumonia by a human commensal *Streptococcus mitis* strain"

**Extended Data Fig. 1**

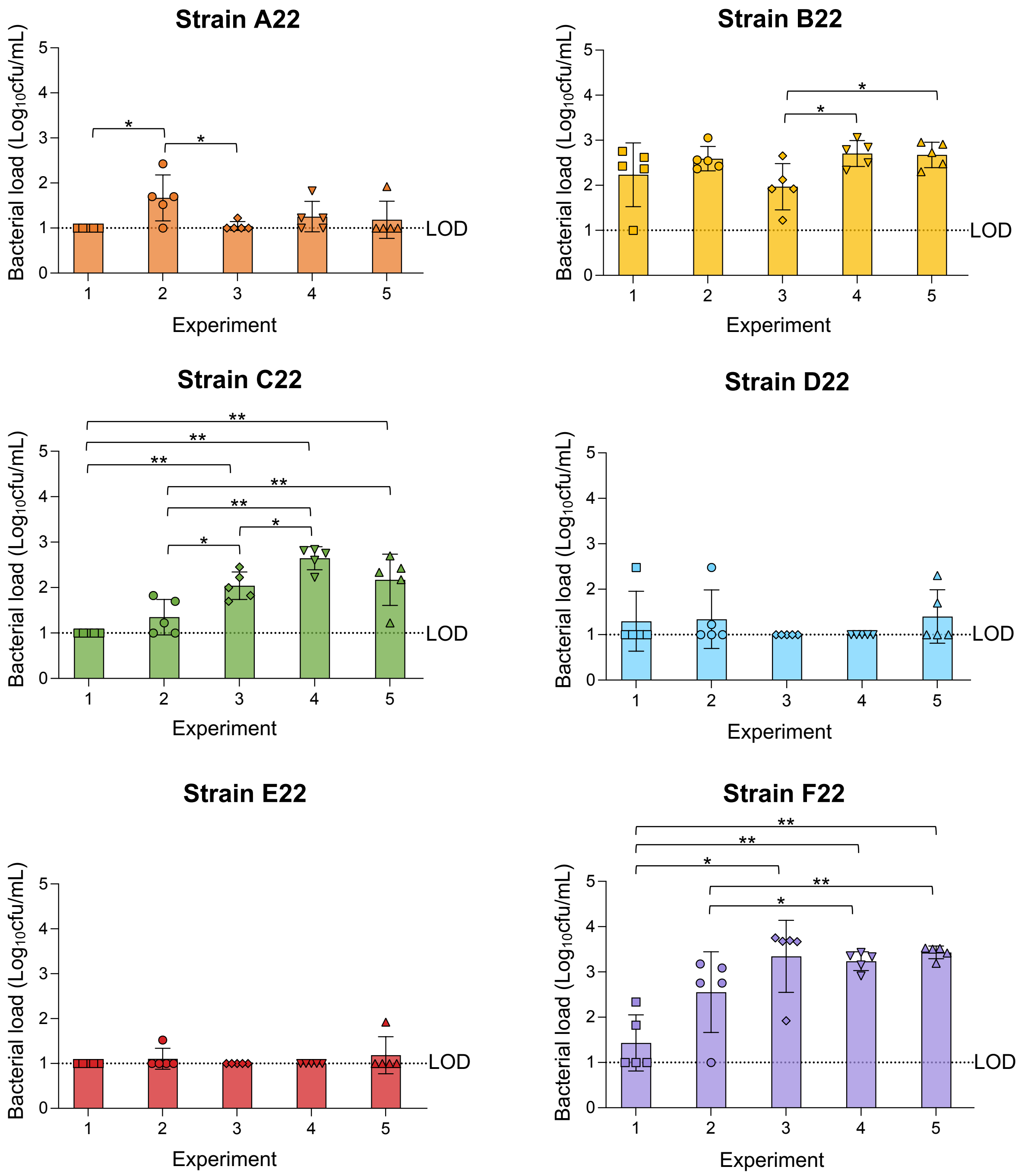


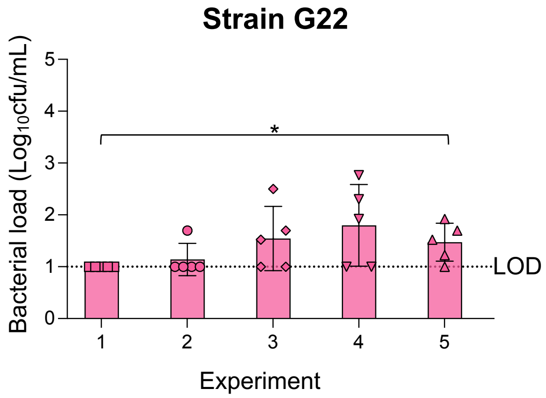


**Extended Data Fig. 1. Bacterial loads obtained from mice colonized with strains A22 to G22.** The colonization potential of strains A22 to G22 was evaluated in different experiments. Bacterial loads were obtained from the nasal lavages three days after inoculation. **Experiment 1:** mice were inoculated with $10^7$ CFU in 10 µL. **Experiment 2:** mice were inoculated with $10^8$ CFU in 10 µL. **Experiments 3 to 5:** Sequential passages were performed. Briefly, the sample from the best colonized mouse of each strain was chosen and inoculated onto a new group of five mice, with minimal laboratory passage, until stabilization of the bacterial density. For strain F22, the best colonizer, the four variants (named F22-1, F22-2, F22-3, and $F22^{Ad}$ from experiments 2, 3, 4, and 5, respectively) were whole-genome sequenced. $^*p < 0.05$, $^{**}p < 0.01$ (Mann-Whitney U test). The dotted line represents the limit of detection (LOD).

**Fig. 2**

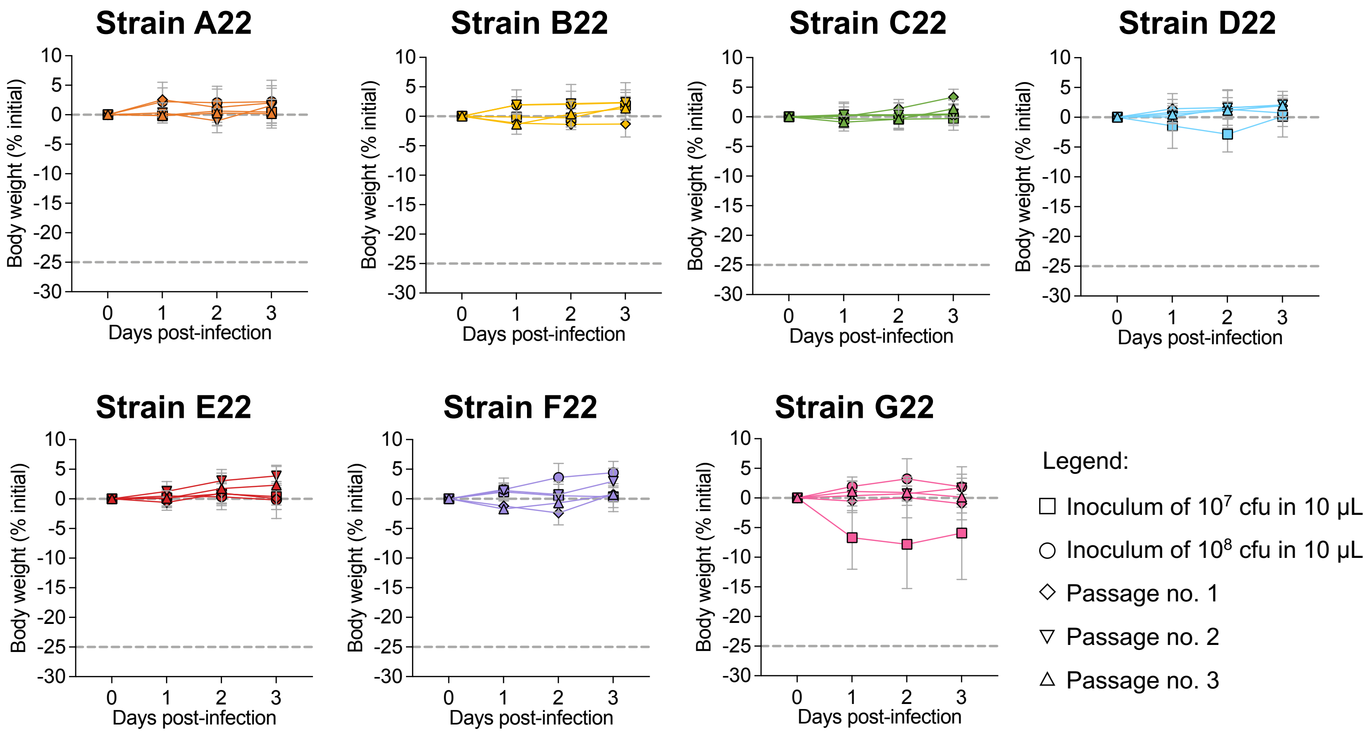


**Extended Data Fig. 2. Weight monitoring of mice during optimization of nasopharyngeal colonization of strains A22 to G22.** Mice were monitored everyday and weight measured. All graphs represent the mean body weight difference, in percentage, of all animals each day post infection.

**Fig. 3**

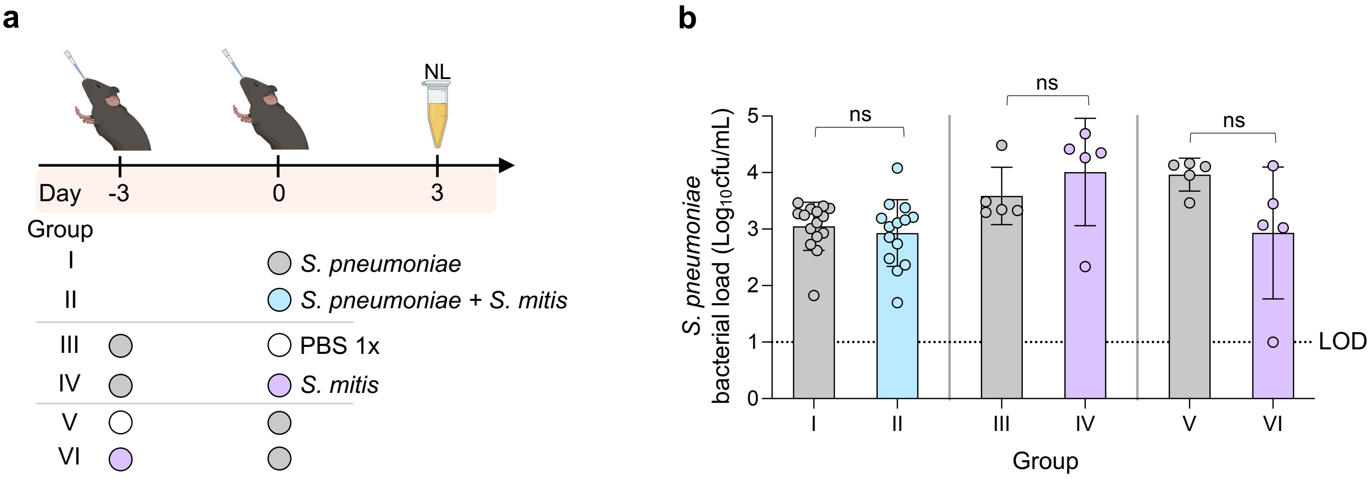


**Extended Data Fig. 3. *In vivo* interaction between *S. pneumoniae* and *S. mitis*. a,** Scheme of the experimental design for the different conditions of the competition assays for *S. pneumoniae* and *S. mitis*. *S. pneumoniae* was inoculated at $10^5$ CFU and *S. mitis* at $10^8$ CFU in 10 µL. Competition was assessed in different timepoints of inoculation: *S. pneumoniae* and *S. mitis* inoculated simultaneously, and bacterial loads assessed after 3 days of colonization (Group II); *S. pneumoniae* inoculated 3 days before or after *S. mitis,* and bacterial loads assessed 3 days after the last inoculation (Groups IV and VI, respectively). Control groups with *S. pneumoniae* inoculated alone or with PBS 1x were performed for each timepoint (Groups I, III and V). Created with BioRender. **b,** Three days after the last inoculation, nasal lavages (NL) were collected and inoculated in parallel in GBA and GBA supplemented with 4 µg/mL of chloramphenicol for strain differentiation. The bacterial loads detected in chloramphenicol plates, indicative of *S. pneumoniae* growth, obtained for each group was plotted. The dotted line represents the limit of detection (LOD). ns: non-significant (Mann-Whitney U test).

**Fig. 4**

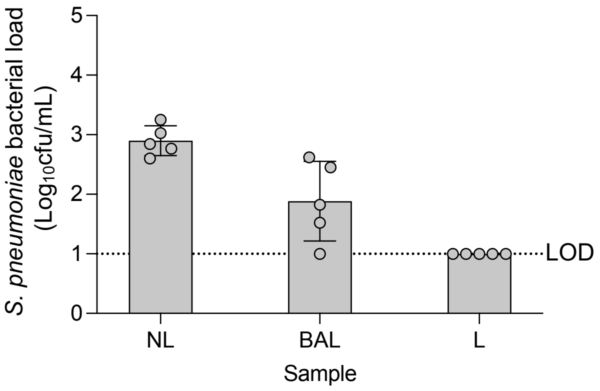


**Extended Data Fig. 4. Bacterial loads of *S. pneumoniae* D39-Cam$^r$ without a previous IAV infection.** Mice were intranasally inoculated with $10^5$ CFU in 10 µL and sacrificed three days after inoculation. Nasal lavages (NL), bronchoalveolar lavages (BAL) and lungs (L) were collected, serial diluted and plated onto gentamicin supplemented blood agar plates. CFUs were determined after incubation. The dotted line represents the limit of detection (LOD).

Fig. 5

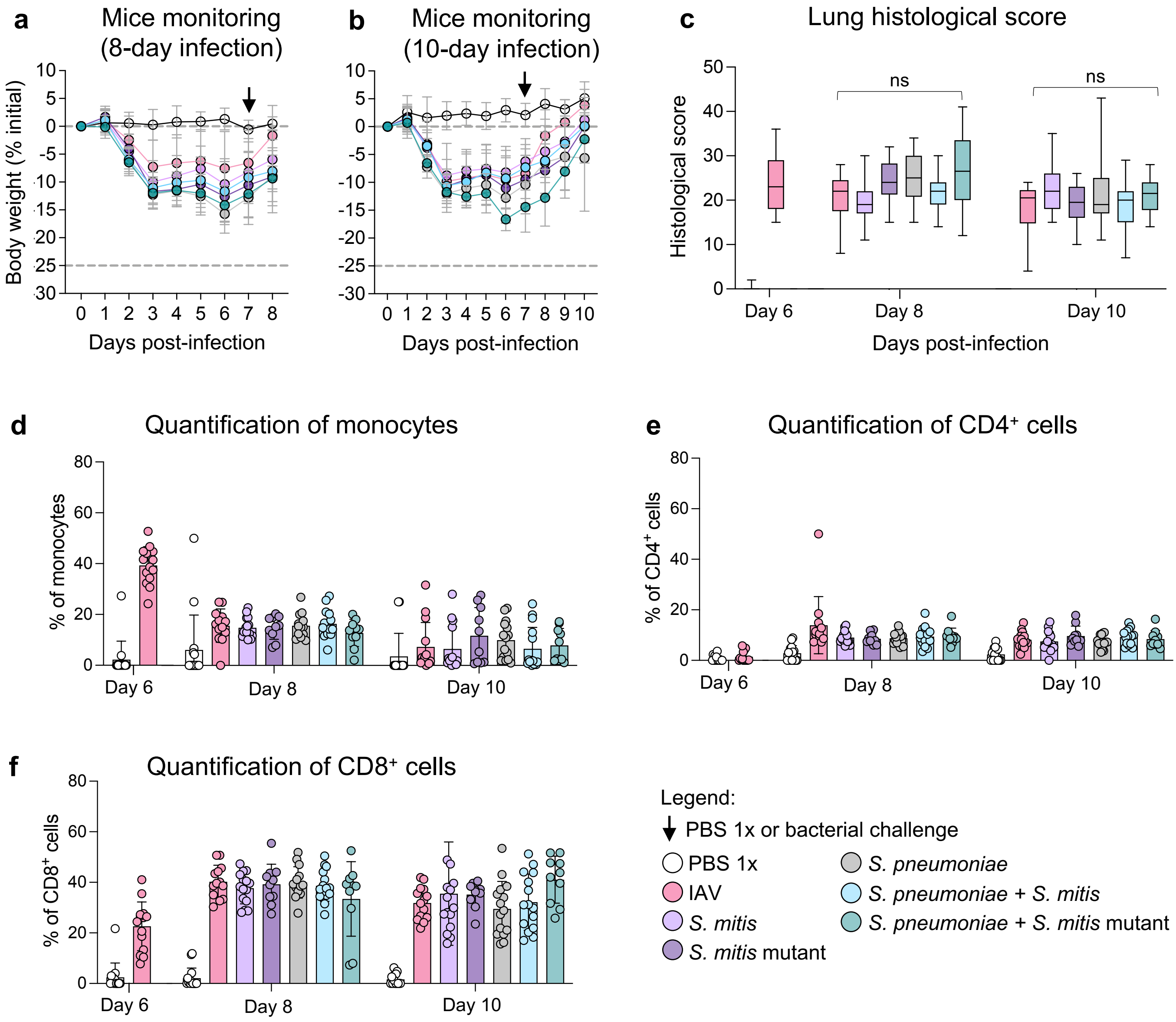


**Extended Data Fig. 5. Infection outcomes in mice. a, b,** The body weight of mice was monitored daily, and the percentage of weight lost or gained was plotted until 8 **(a)** or 10 **(b)** days post-IAV infection, time at which the animals were sacrificed. The black arrow indicates the time-point of bacterial inoculation, by day 7 post-IAV infection. **c,** Histological score of lungs in terms of damage and inflammation was performed blindly by a pathologist. **d, e, f,** Flow cytometry of BAL samples was performed for all groups and the presence of monocytes **(d)**, $CD4^+$ **(e)** and $CD8^+$ T cells **(f)** was analysed.

Fig. 6

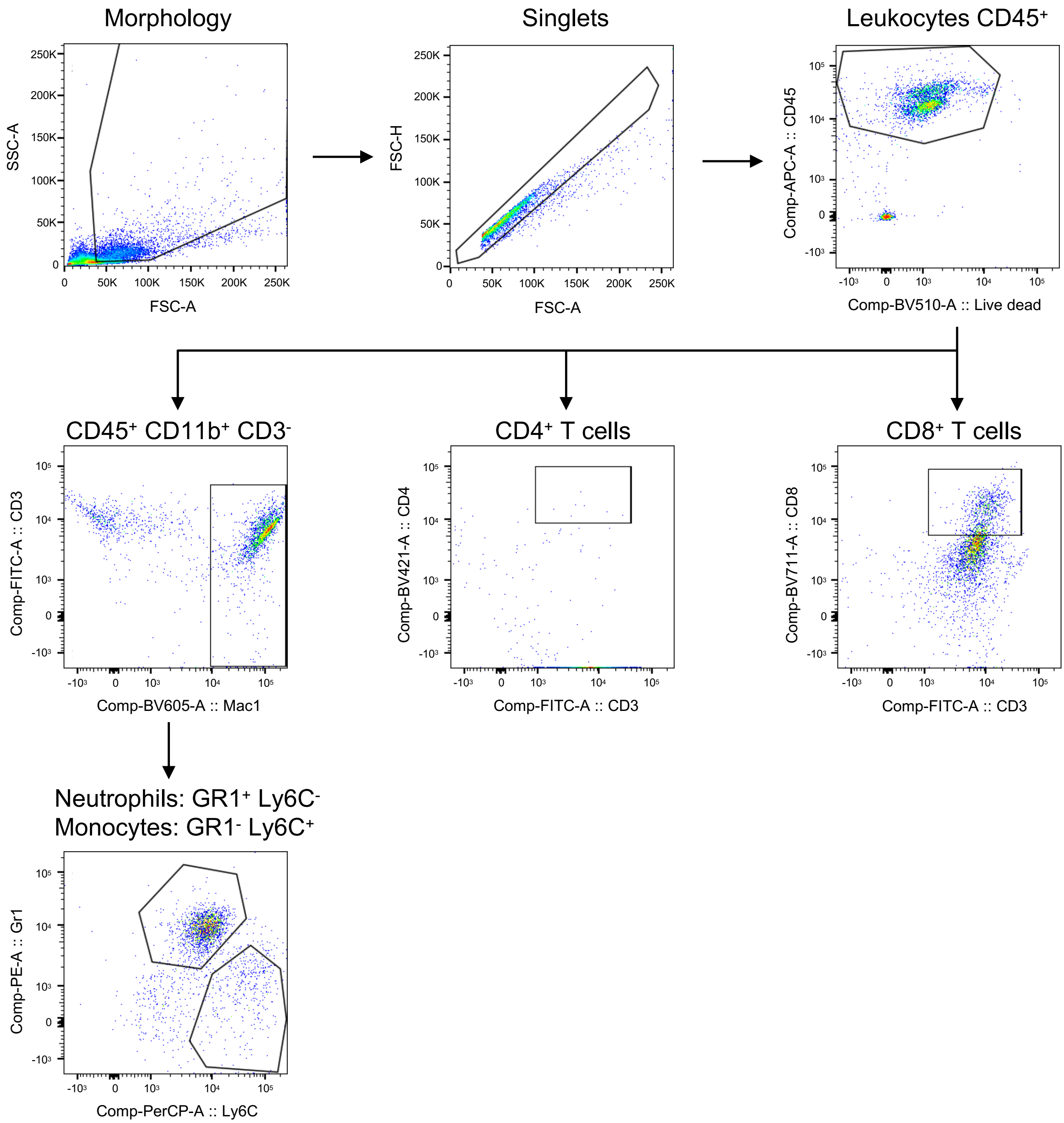


**Extended Data Fig. 6. Gating strategy for flow cytometry of BAL samples.** Flow cytometry analysis of cell populations was performed in a BD LSR Fortessa X-20 SORP (BD Biosciences) equipped with BD FACSDiva™ 8 (BD Biosciences) and FlowJo™ software v10.10 (BD Life Sciences). Cell populations were first gated by forward and side scatter to exclude debris, followed by doublet discrimination using FSC-A versus FSC-H. Live single cells were then gated as $CD45^+$, and immune subsets were identified based on expression of specific markers: monocytes ($CD11b^+$ $Ly6C^+$), neutrophils ($CD11b^+$ $GR1/Ly6G^+$), $CD4^+$ T cells ($CD3^+$ $CD4^+$), and $CD8^+$ T cells ($CD3^+$ $CD8^+$).

### Fig. 7

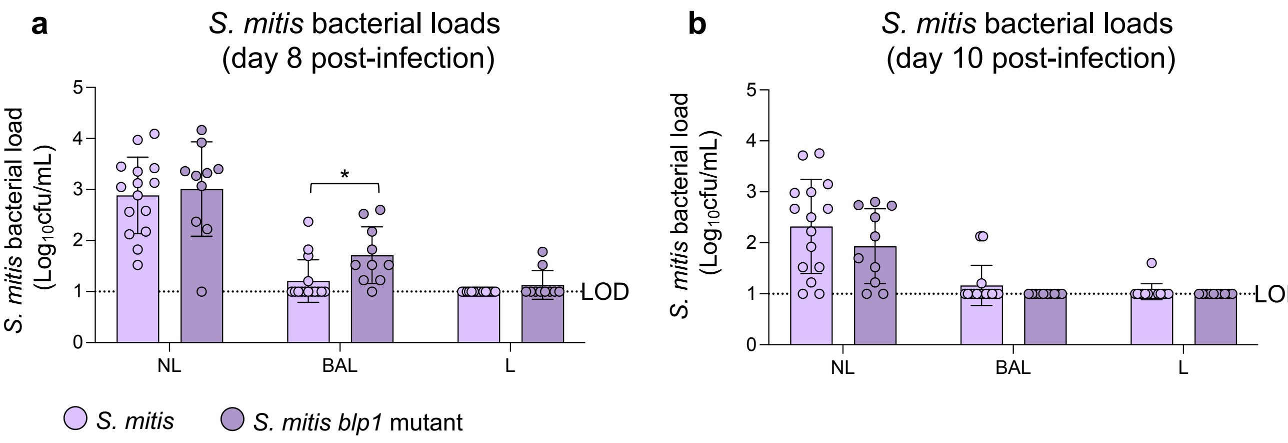


**Extended Data Fig. 7. Bacterial loads of *S. mitis* F22$^{Ad}$ and F22$^{Ad}$Δ*blp1* in the context of an IAV infection.** Seven days after IAV infection, mice were intranasally inoculated with $10^7$ CFU in 10 µL of F22$^{Ad}$ or F22$^{Ad}$Δ*blp1*. Eight **(a)** or ten-days **(b)** after inoculation, nasal lavages (NL), bronchoalveolar lavages (BAL) and lungs (L) were collected, serial diluted and plated onto gentamicin supplemented blood agar plates. CFUs were determined after incubation. The dotted line represents the limit of detection (LOD). *p < 0.05 ( Mann-Whitney U test to compare two groups)

### Fig. 8

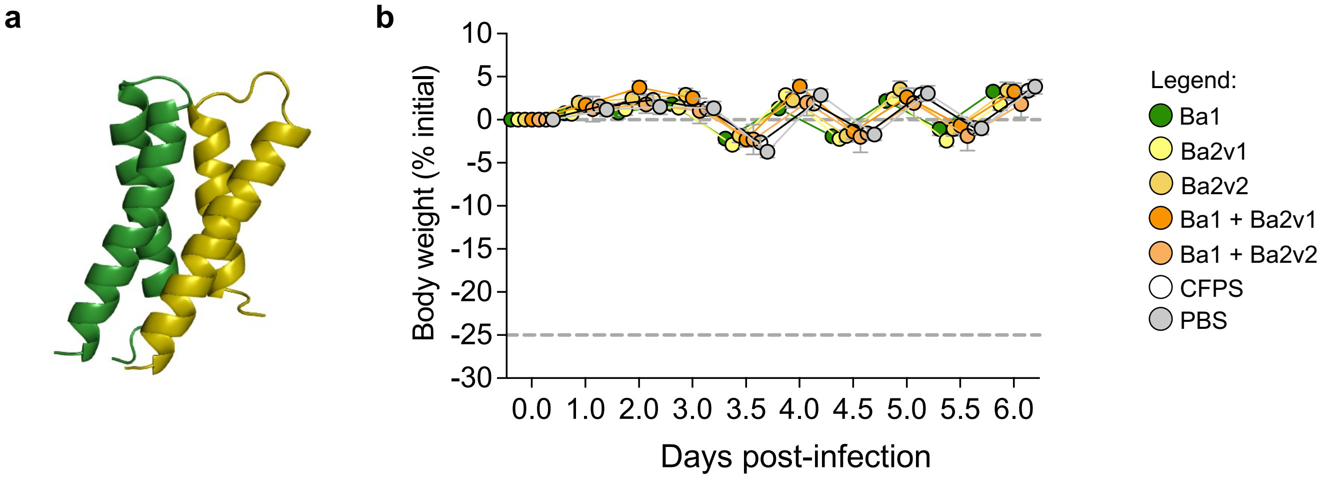


**Extended Data Fig. 8. Effect of bacteriocins on pneumococcal colonization. a,** AlphaFold prediction of the two-peptide structure of Bac1 (green) and Bac2v1 (yellow) that exhibited and inhibitory activity towards *S. pneumoniae* colonization. **b,** Weight monitoring of mice during treatment with all bacteriocins from *S. mitis* strain F22 and their combinations, after three-day colonization with *S. pneumoniae* D39-Cam$^r$. Mice were monitored everyday and weight measured. All graphs represent the mean body weight difference, in percentage, of all animals each day post infection.

**Fig. 9**

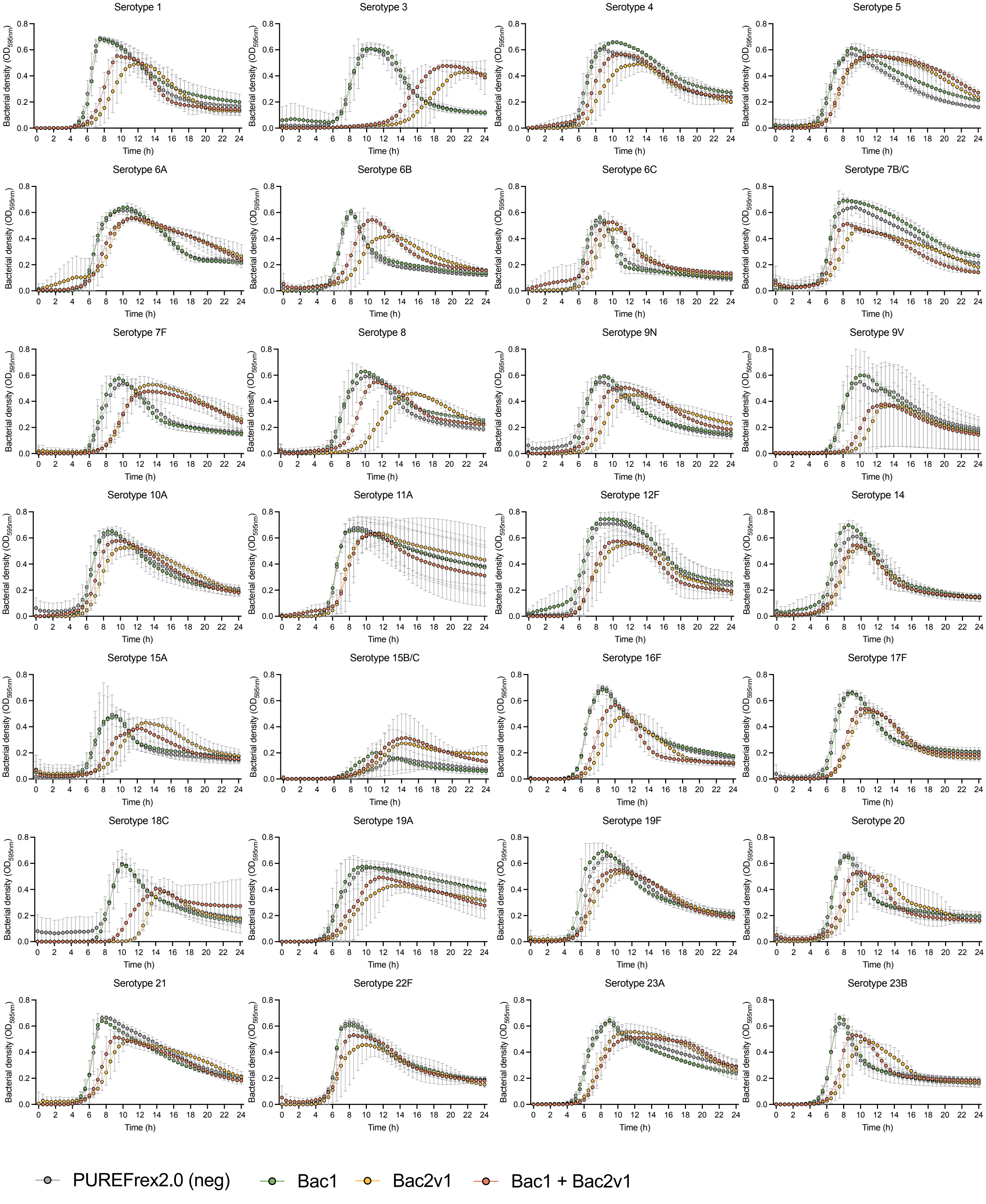


(continues next page)

**Fig. 9 (cont.)**

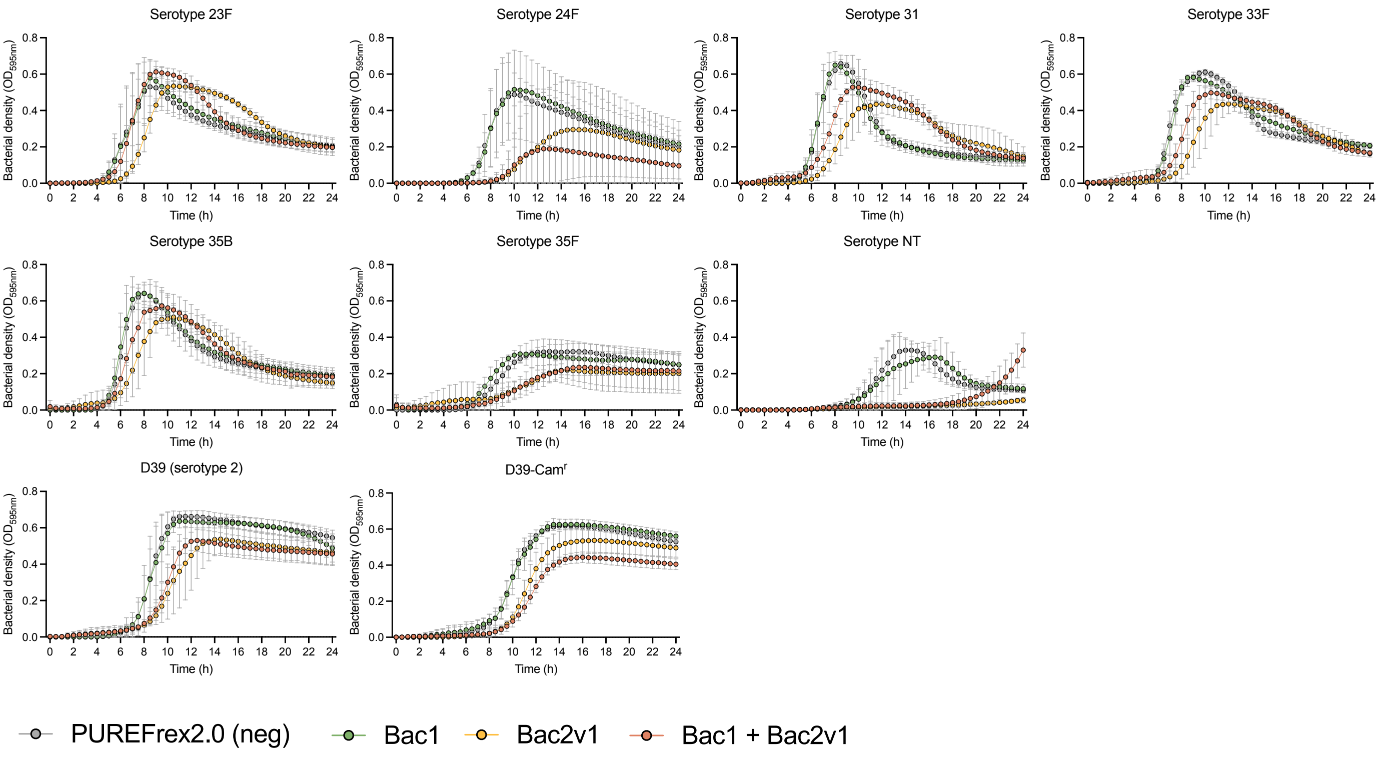


**Extended Data Fig. 9. Effect of bacteriocins on 36 epidemiologically relevant *S. pneumoniae* serotypes.** Cell-free synthesized bacteriocins were tested against 36 serotypes of *S. pneumoniae***.** Pneumococcal cultures were grown in C+$Y_{YB}$ medium at 37ºC until reaching an $OD_{600nm}$ of 0.5. These cultures were then diluted to $10^4$ CFU/mL in fresh C+$Y_{YB}$ medium, distributed in 384-well plates (90 µL per well) and treated with 10 µL of cell-free synthesized bacteriocin (when in single; 5 µL of each when in combination). Controls were treated with the same volume of the CFPS reaction using water instead of DNA. Growth was monitored for 24 hours by measuring $OD_{595nm}$ every 30 minutes using a plate reader (Tecan Infinite 200 Pro). Three independent experiments were performed for each bacteriocin.
